# DeepBacs: Bacterial image analysis using open-source deep learning approaches

**DOI:** 10.1101/2021.11.03.467152

**Authors:** Christoph Spahn, Romain F. Laine, Pedro Matos Pereira, Estibaliz Gómez-de-Mariscal, Lucas von Chamier, Mia Conduit, Mariana Gomes de Pinho, Guillaume Jacquemet, Séamus Holden, Mike Heilemann, Ricardo Henriques

## Abstract

Deep Learning (DL) is rapidly changing the field of microscopy, allowing for efficient analysis of complex data while often out-performing classical algorithms. This revolution has led to a significant effort to create user-friendly tools allowing biomedical researchers with little background in computer sciences to use this technology effectively. Thus far, these approaches have mainly focused on analysing microscopy images from eukaryotic samples and are still underused in microbiology. In this work, we demonstrate how to use a range of state-of-the-art artificial neural-networks particularly suited for the analysis of bacterial microscopy images, using our recently developed ZeroCostDL4Mic platform. We showcase different DL approaches for segmenting bright field and fluorescence images of different bacterial species, use object detection to classify different growth stages in time-lapse imaging data, and carry out DL-assisted phenotypic profiling of antibiotic-treated cells. To also demonstrate the DL capacity to enhance low-phototoxicity live-cell microscopy, we showcase how image denoising can allow researchers to attain high-fidelity data in faster and longer imaging. Finally, artificial labelling of cell membranes and predictions of super-resolution images allow for accurate mapping of cell shape and intracellular targets. To aid in the training of novice users, we provide a purposefully-built database of training and testing data, enabling bacteriologists to quickly explore how to analyse their data through DL. We hope this lays a fertile ground for the efficient application of DL in microbiology and fosters the creation of novel tools for bacterial cell biology and antibiotic research.

## Introduction

The study of microorganisms and microbial communities is a multidisciplinary approach bringing together molecular biology, biochemistry, and biophysics. It covers large spatial scales ranging from single molecules over individual cells to entire ecosystems. The amount of data collected in microbial studies constantly increases with technical developments, which can become challenging for classical data analysis and interpretation, requiring more complex computational approaches to extract relevant features from the data landscape. Therefore, manual analysis is replaced increasingly by automated analysis, notably with machine learning (ML) (1). In bioimage analysis, ML contributed to a better understanding of viral organisation (2) and the mode of action of antimicrobial compounds (3). In recent years, the interest in deep learning (DL) tools for bioimage analysis has significantly increased, as their high versatility allows them to perform many different image analysis tasks with high performance and speed (4–7). This was impressively demonstrated for image segmentation (8–12), artificial labelling (13, 14) or recovery of high-quality (15, 16) and even prediction of super-resolution images (15, 17, 18) from low-quality images. Other networks facilitate image-to-image translation (19) or object detection and classification (20, 21). Next to the development of novel DL approaches, effort has been put into their democratisation and in providing an entry-point for non-experts by simplifying their use and providing pretrained models (22–25). To further democratise expensive model training, recent developments employ cloud-based hardware solutions, thus bypassing the need for specialised hardware (23, 26, 27). However, these methodologies dominantly focused on the study of eukaryotes, particularly given the wealth of pre-existing imaging data (18, 28). In image-based microbiology, DL is used mainly for segmentation (29–32), while other DL-assisted bioimage analysis tasks remain largely underexploited. To address this gap, we propose that existing open-source DL approaches can be easily expanded or adapted to analyse bacterial bioimages. As the key requirement for successful application of DL is suitable training data, we generated various image datasets comprising different bacterial species (*E. coli, S. aureus* and *B. subtilis*) and imaging modalities (bright field, widefield and confocal fluorescence imaging and super-resolution microscopy) (Supplementary Table 1). We used these datasets to train DL models for a wide range of applications (Figure 1) using the recently developed Zero-CostDL4Mic platform (23), benefitting from its ease-of-use and low-cost capabilities. Specifically, we demonstrate the potential of open-source DL technology in image segmentation of both rod and spherically shaped bacteria (both fluorescence and noisy bright field images); in the detection of cells and their classification based on growth stage and antibiotic-induced phenotypic alterations; on denoising of live-cell microscopy data, such as nucleoid and FtsZ dynamics; and finally, we explore the potential of DL approaches for artificial labelling of bacterial membranes in bright field images and prediction of super-resolution images from diffraction-limited widefield images. We share our data and models for researchers to explore the different DL networks and to test and use pretrained models on existing data. Using pretrained models, researchers can train custom models more efficiently via transfer learning, requiring less time and less data to reach high performance (23, 33). We envision that this work will help microbiology researchers seamlessly leverage DL for microscopy image analysis, allowing them to benefit from high performance and high-speed algorithms.

**Fig. 1.**
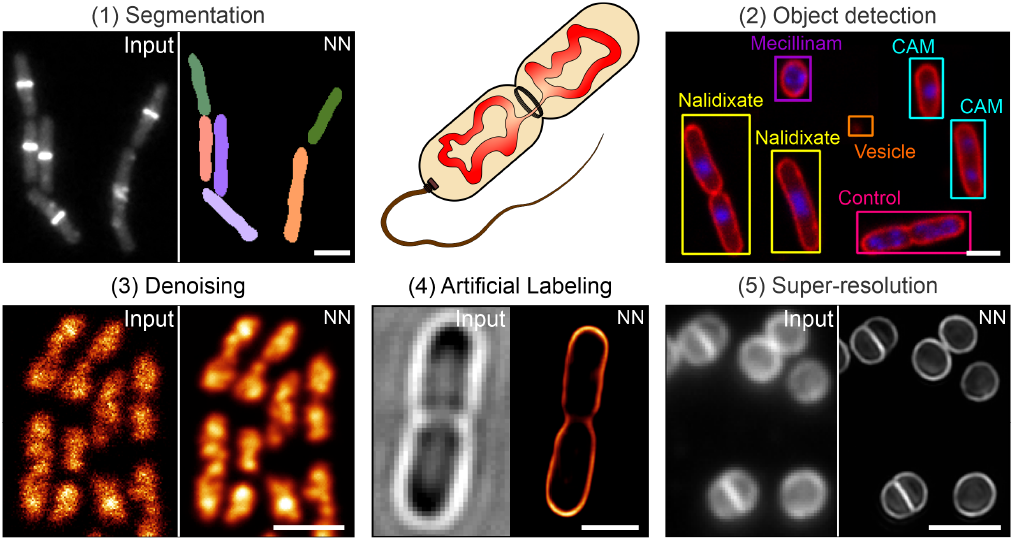
Overview of the DL tasks and datasets used in DeepBacs. We demonstrated the capabilities of DL in microbiology for segmentation (1), object detection (2), denoising (3), artificial labelling (4) and prediction of super-resolution images (5) of microbial microscopy data. A list of datasets can be found in Table S1, comprising different strains such as *B. subtilis* (1), *E. coli* (2-4) and *S. aureus* (5) and imaging modalities (i.e. widefield and confocal fluorescence microscopy, bright field imaging or super-resolution techniques). NN: neural network output. CAM = Chloramphenicol. Scale bars: 2 µm.

## Results

In the following sections, we describe the individual datasets (Supplementary Table 1) that we used to perform the tasks shown in Figure 1. We explain how we designed our experiments, how data was prepared and analysed, and showcase results for the different image analysis tasks. Datasets and selected trained models are publicly available via the data-sharing platform Zenodo (Supplementary Table 2), allowing users to explore the DL technology described in this work.

### Image segmentation

Up to date, image segmentation represents the main application of DL technology for bacterial bioimages. It facilitates single-cell analysis in larger image analysis pipelines and automated analysis of large datasets (29–32, 34, 35). Due to the considerable variety in microscopy techniques and bacterial shapes, to our knowledge, there is no universal DL network that excels for all types of data, leading to the emergence of many highly specific networks. However, the requirement of specific hardware prevents microbiologists from applying them to their data. We thus sought to test different easily applicable DL networks for their capability to segment the types of bacterial bioimages frequently encountered in microbiological studies. For this, we generated and annotated different datasets (see methods), comprising bright field and fluorescence microscopy images of rod- and cocci-shaped bacteria (*E. coli* and *S. aureus* for bright field, *S. aureus* and *B. subtilis* for fluorescence) (Figure 2). For all datasets, we trained DL models using ZeroCostDL4Mic, as it provides simple and rapid access to a range of popular DL networks (23). To evaluate their performance, we calculated common metrics, which compare the network output of test images to the respective annotated ground truth (see Supplementary Note 1). Here we discriminate between semantic and instance segmentation. Semantic segmentation mostly creates a binary image of the background and foreground pixels, while instance segmentation extracts individual objects from semantic segmentations. Model performance in semantic segmentation is assessed using the intersection-over-union (IoU) metric that reports on the overlap of output and ground truth segmentation masks, with higher overlap representing better agreement. To assess the quality of instance segmentation, we determined the recall and precision metrics, which report on the model sensitivity and specificity (Supplementary Note 1).

**Fig. 2.**
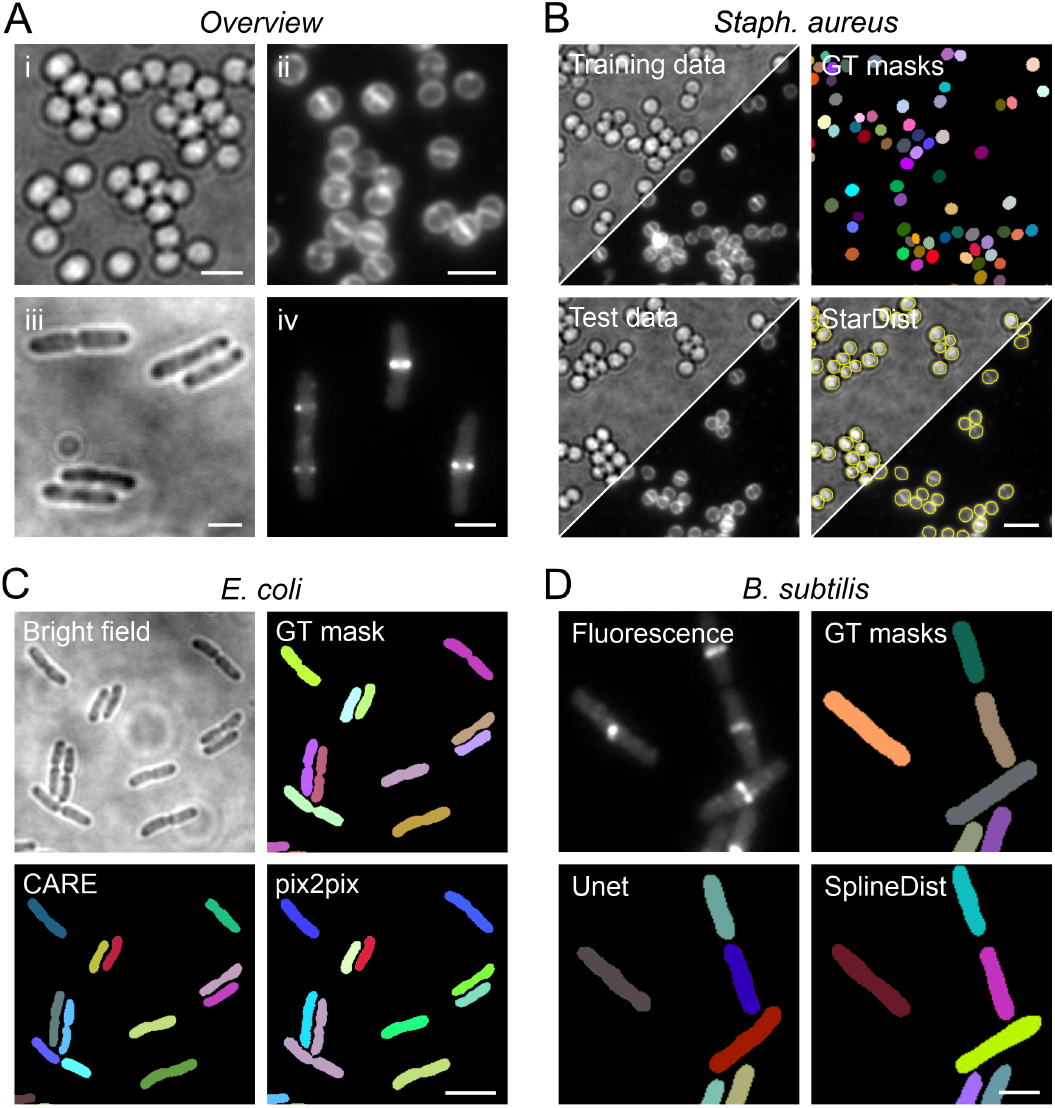
Segmentation of bacterial images using open-source deep learning approaches. **[A]** Overview of the datasets used for image segmentation. Shown are representative regions of interest for (i) *S. aureus* bright field and (ii) fluorescence images (Nile Red membrane stain), *E. coli* bright field images (iii) and (iv) fluorescence images of *B. subtilis* expressing FtsZ-GFP. **[B]** Segmentation of *S. aureus* bright field and membrane-stain fluorescence images using StarDist (9). Bright field and fluorescence images were acquired in the same measurements. Yellow dashed lines indicate the cell outlines detected in the test images shown. **[C]** Segmentation of *E. coli* bright field images using the U-Net type network CARE (15) and GAN-type network pix2pix (19). A representative region of a training image pair (bright field and GT mask) is shown. **[D]** Segmentation of fluorescence images of *B. subtilis* expressing FtsZ-GFP using U-Net and SplineDist (36). GT = ground truth. Scale bars are 2 µm (C, E) and 3 µm (B, D).

Five popular networks were used for segmentation, namely U-Net (8), CARE (15), pix2pix (19), StarDist (9) and its recent variant SplineDist (36) (Supplementary Table 3). As the underlying network architectures vary, the workflows to obtain instance segmentations differ in terms of e.g. input/output data formats and image post-processing (Figure S1). While StarDist and SplineDist provide instance segmentation directly as network output, instances have to be generated from U-Net, CARE and pix2pix predictions by post-processing the outputs. CARE and pix2pix were not explicitly designed for segmentation but are versatile enough to generate segmentable probability maps. Similar to U-Net, the instance segmentation performance does not only depend on the trained model, but also depends on the applied post-processing routine.

For our first dataset, we recorded live *S. aureus* cells immobilised under agarose pads, either in bright field mode or using the fluorescent membrane stain Nile Red (Figure 2B). Due to their coccoid shape, we speculated that StarDist (9) is well suited to segment this kind of data. StarDist is a U-Net-based network developed to segment star-convex objects (e.g. cell nuclei) at high object densities. We manually annotated 7 images with varying cell densities (46 - 335 cells) and split these into 28 image patches. We then trained two independent StarDist models (one for bright field and one for fluorescence image segmentation) using the ZeroCostDL4Mic StarDist implementation. As a general strategy to increase the training dataset, we used data augmentation (23, 37) for all DL learning tasks performed in this study. Testing an unseen and fully annotated dataset demonstrated cell counting accuracies (recall) of 98 ± 2% (membrane fluorescence) and 90 ± 2% (bright field). The reduced accuracy for bright field images is caused by optical artefacts at high cell density, leading to merging of defocussed cells (see Figure 2Ai). Next to performing segmentation in the cloud, trained models can also be downloaded and conveniently used with the StarDist plugin distributed via the image analysis platform Fiji (38) (Supplementary Video 1). Similar to the Zero-CostDL4Mic notebook, this enables efficient segmentation of live-cell time-lapse data (Supplementary Video 2).

Motivated by this finding, we sought to know whether StarDist is also suitable to segment rod-shaped cells. For this, we recorded bright field time-lapse images of live *E. coli* cells immobilised under agarose pads (39) (Figure 2C). Bright field images show less contrast than phase contrast images and suffer from high noise, making them challenging to be segmented. Still, they are widespread in bacterial imaging, and proper segmentation would be beneficial to study bacterial proliferation (i.e. cell counts over time) and morphology (cell dimensions and shape). We annotated individual image frames spread over the entire time series and trained supervised DL networks to reflect varying cell sizes and densities. All networks showed a good performance for semantic segmentation, as indicated by high IoU values for all time points (IoU > 0.75) (Supplementary Figure 2A). For instance segmentation, however, the model performance varied strongly. We found that instance segmentation of U-Net, CARE and pix2pix worked well for early time points and thus low cell density (Figure 2B). Individual cells in crowded regions could not be resolved using basic image post-processing (i.e. thresholding), which led to a successive decrease of the recall value over time and thus a decreasing number of correctly identified cells (Supplementary Figure 2A). The best counting performance at low and high cell densities was achieved using StarDist, which correctly identified 87% of the cells for the entire test dataset. However, as StarDist assumes star-convex shaped objects, the accuracy of the predicted cell shape decreased with increasing cell length (and thus axial ratio), rendering this network less suited for morphometry of elongated rod-shaped cells (Supplementary Figure 2B). Using a multilabel U-Net (trained to detect cell cytosol and boundary) instead of a single-label U-Net provided the best compromise between instance segmentation performance and proper prediction of cell morphology. Applying this model to time-lapse videos allows to extract single-cell instances that can subsequently be tracked using e.g. TrackMate (40) (Supplementary Video 3).

Next, we were interested in the performance of DL networks for the segmentation of complex fluorescence data. While typical images used for segmentation show a high contrast (fluorescent membrane stains, phase contrast images), images with complex fluorescence distributions or low signal represent a significant challenge. This is particularly true for classical segmentation methods like intensity thresholding. We here demonstrate this task being applied on fluorescence images of *B. subtilis* cells expressing FtsZ-GFP (41). Next to the characteristic localisation in the septal region, diffusing FtsZ monomers produce dim labelling of the cytosol, leading to a heterogeneous intensity distribution (Figure 2D). Growth for several cell cycles results in the wellknown *B. subtilis* filaments and microcolonies, providing a dataset with increasing cell density and a large amount of cell-to-cell contacts. When we tested different networks for this challenging dataset, we found that U-Net and pix2pix provided well segmentable predictions at low to medium cell density (Figure 2D, Supplementary Figure 2C/D). At high cell densities, however, these networks also suffered from the undesired merging of cells, leading to reduced recall and precision values (Supplementary Figure 2C/D). As for segmentation of *E. coli* bright field images, StarDist and its variant SplineDist (36) showed high recall and precision values also for mid- and high-density regions, while the multi-label U-Net preserved cell morphology at slightly lower instance segmentation accuracy (Supplementary Figure 2D). In contrast to StarDist, SplineDist is not limited to convex shapes, which makes it a good candidate network for the segmentation of curved bacteria (e.g. *Caulobacter crescentus*). It is, however, computationally more expensive and thus supports smaller training datasets than StarDist using the Zero-CostDL4Mic platform.

Finally, and motivated by generalist approaches such as Cellpose (11), we were interested in whether a single StarDist model is able to perform all the segmentation tasks presented in this study. We thus trained a model on pooled training data and evaluated its performance (Supplementary Figure 3). Although bright field and fluorescence images differ significantly, the obtained ‘all-in-one’ model showed similar precision and recall values compared to the specialist models (Supplementary Table 4). However, it also shows the same limitations, such as incomplete predictions for long and curved bacteria (Supplementary Figure 3, lowest panel). For cells with suitable morphology, StarDist allows segmenting large images with thousands of cells, as shown for live *Agrobacterium tumefaciens* imaged at various magnifications (Supplementary Figure 4). It also demonstrates that DL segmentation models can perform well on images with low signal and noisy background.

### Object detection and classification

Object detection is a task closely related to image segmentation. However, instead of the network classifying pixels as background or foreground pixels, it outputs a bounding box and class label for each detected object. This is used extensively in real-life applications, such as self-driving cars or detecting items in photographs (20). To explore the potential of object detection for microbiological applications, we employed an implementation of YOLOv2 (20) for two distinct tasks: Identification of cell cycle events such as cell division in bright field images (Figure 3A) and antibiotic phenotyping using membrane and DNA stains (Nile Red and DAPI, respectively) (Figure 3B) (Supplementary Table 5). These labels are commonly used to study antibiotic action, as they are easy to use and also facilitate live-cell staining of bacterial cells (42). We chose YOLOv2 due to its good performance in a recent study, in which a network was trained to classify cell nuclei in fluorescence images (43). For growth stage classification of live *E. coli* cells in bright field images we used the same dataset employed for segmentation (Figure 2C). Here, we wanted to discriminate between rod-shaped cells, dividing cells, and microcolonies (Figure 3A). Microcolonies emerge during constricted growth under agarose pads, but can also be caused by cell clumping under unfavourable growth conditions. Imaging such regions is often undesired, as they complicate single-cell studies, and automated skipping can thus save time and resources. To test YOLOv2 for growth stage classification we annotated a set of bright field images by drawing bounding boxes around nondividing cells, dividing cells and microcolonies (4+ cells in close contact) online (https://www.makesense.ai/) or locally (LabelImg) (44) (see methods).

**Figure 3:**
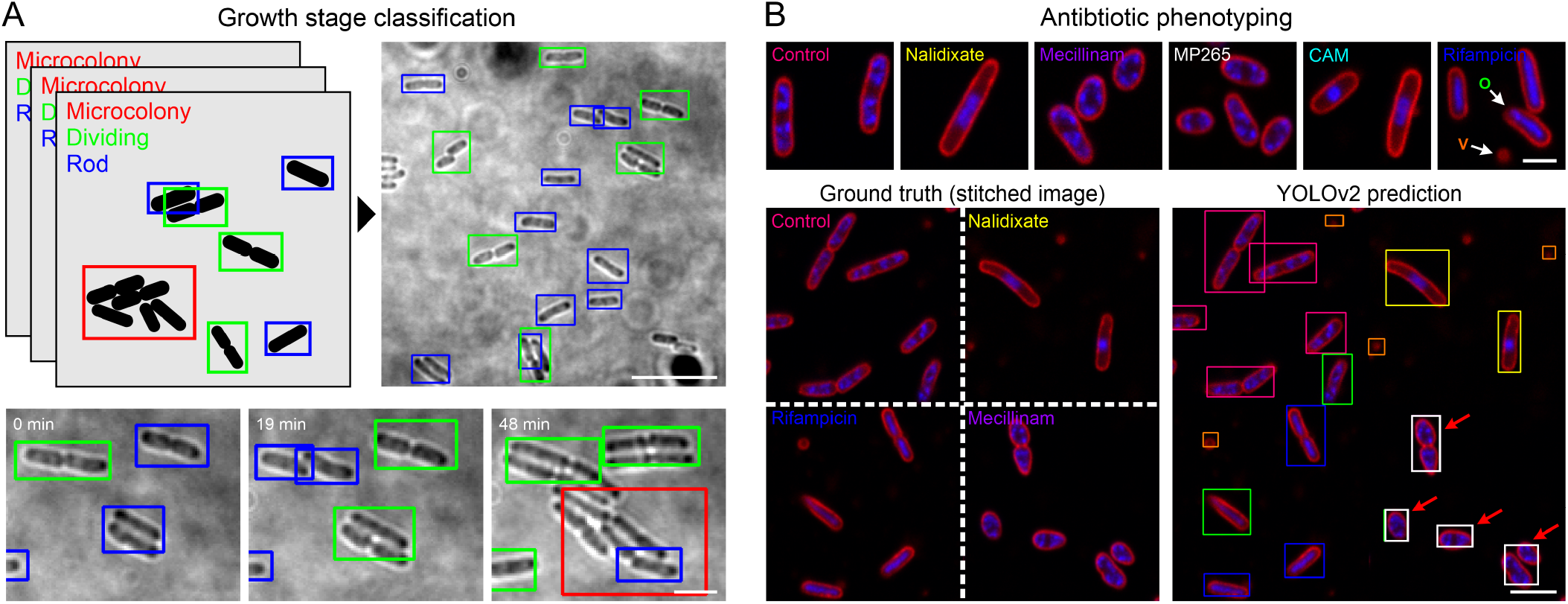
DL-based object detection. **[A]** A YOLOv2 model was trained to detect and classify different growth stages of live *E. coli* cells. “Dividing” cells (green bounding boxes) show visible septation, the class “Rod” (blue bounding boxes) represents growing cells without visible septation and regions with high cell densities are classified as “Microcolonies” (red bounding boxes). Lower panel shows three frames from a live cell measurement. **[B]** Antibiotic phenotyping using object detection. A YOLOv2 model was trained on drug-treated cells (upper panel). The model was tested on synthetic images randomly stitched from patches of different drug treatments. Bounding box colors refer to the color-code in the upper image panel, vesicles (V, orange boxes) and oblique cells (O, green boxes) were added as additional classes during training. Mecillinam-treated cells were misclassified as MP265-treated cells (red arrows). Scale bars are 10 µm (A, overview), 3 µm (B, lower panel) and 1 µm (B, upper panel).

Initially, we wanted to know how the object size influences the performance of YOLOv2. We thus annotated both whole images (512 × 512 px^2^) and cropped images (256× 256 px^2^). Since object detection networks scale input images to a specific size (for YOLOv2: 416 × 416 px^2^), cropping results in a larger relative object size in the network input. When we trained our model on large images, we encountered missed objects, wrong bounding box positioning or false classifications (Supplementary Video 4). To quantify this effect, we applied the model to the annotated test dataset and determined recall and precision values as well as the mean average precision (mAP, see Supplementary Note 1) (Supplementary Table 6). mAP represents the common metric for object detection, taking into account model precision and recall over a range of object detection thresholds (20). For object detection challenges (e.g. PASCAL visual object classes (VOC) challenge (45)), well-performing models typically yield mAP values in the range of 0.6 - 0.8. However, for our large-FoV growth stage classification dataset, we obtained a mAP of 0.386. The observed per-class AP values hereby show the size-dependent performance in object detection (AP_Microcolony_ > AP_dividing_ > AP_rod (non-dividing)_) (Supplementary Table 6). Using cropped images resulted in improved network performance on the classification of most cells in the image (Figure 3A) and an improved mAP of 0.667 (Supplementary Table 6). Knowing the size-dependent performance is important for the design of an object detection experiment. If the focus is e.g on the detection of large structures, e.g. microcolonies, the YOLOv2 model can be trained on large fields of view. However, small regions of interest or higher optical magnification should be used if small objects are to be detected. Object density is another parameter that affects performance of YOLOv2 models. To identify objects in different parts of the image, YOLOv2 divides the image into a grid, and each grid region can only contain a single object. For the YOLOv2 implementation that we employed, this is a 13×13 grid, resulting in a maximum of 169 objects detected for optimal object distribution. If the centroids of two or more bounding boxes fall into the same region, only one object will be detected. Very close objects (i.e. non-dividing cells at t = 0 in Figure 3A or dividing cells at t = 19 min, yellow arrows) are hence not resolved, instead only one bounding box of the corresponding class is predicted. Object density should thus be considered as a limiting factor when planning to train a network for object detection. When applied to time-lapse recordings of growing *E. coli* cells, the model facilitates identification of class transitions, e.g. from rod-shaped (non-dividing) cells to dividing cells and at later time points to microcolonies (Figure 3A, lower panel, Supplementary Video 5).

As a second task for object detection, we explored its suitability for antibiotic phenotyping (Figure 3B). In antibiotic phenotyping, bacterial cells are classified as non- or drug-treated cells based on cell morphology and subcellular features (commonly DNA and membrane stains). This facilitates the assignment of a mode of action to antibiotics or potential candidate compounds, being a promising tool in drug discovery (3, 42). To explore whether object detection networks can be used for this purpose, we generated a dataset of images including membrane- and DNA-labelled *E. coli* cells grown in the absence or presence of antibiotics. We used five different antibiotics that target different cellular pathways (Figure 3B). Nalidixate blocks DNA gyrase and stalls DNA replication. Mecillinam and MP265 perturb cell morphology by inhibiting peptidoglycan crosslinking by PBP2 or MreB polymerisation, while rifampicin and chloramphenicol inhibit transcription and translation, respectively (Supplementary Table 7). As additional classes, we included untreated cells (control), membrane vesicles and oblique cells. The latter class represents cells that are only partially attached to the surface during immobilisation. Such cells can be identified by a focus shift and are present in all growth conditions (Figure 3B, upper panel). Further examples for each class are provided in Supplementary Figure 5. We trained a YOLOv2 model on our annotated dataset and tested its performance on two test datasets: The first dataset consists of images containing bacteria with different treatments (stitched images, see methods) (Figure 3B, lower panel). These images were randomly assembled from patches of different drug treatments by merging four 200 × 200 px^2^ patches on a 2 × 2 grid. The second test dataset includes images that only show one condition, similar to the dataset used for model training (Supplementary Figure 4). The YOLOv2 model showed a comparable performance for both datasets with mAP values of 0.66 (stitched image dataset) and 0.69 (individual conditions), indicating a good generalisability of our model. Interestingly, AP values for the different classes varied substantially, ranging from 0.21 (vesicles) to 0.94 (control) (Supplementary Table 8). Poor prediction of membrane vesicles is likely caused by their small size, which agrees with the observations made for growth stage prediction. Intermediate AP values are observed when antibiotics induce similar morphological changes, as it is the case for mecillinam (AP = 0.605) and MP265 (AP = 0.526). This led to misclassification between these classes (Figure 3B, red arrows, Supplementary Figure 6), indicating that both treatments result in a highly similar phenotype. This similarity allowed us to test whether YOLOv2 can identify antibiotic modes of actions in unseen images. We omitted MP265 data during model training, but included images of MP265-treated cells in the test data. Due to their similar phenotype MP265-treated cells should hence be predicted as Mecillinam-treated. This was indeed the case, as shown by the high mAP value (0.866) and specificity (recall = 0.961) (Supplementary Figure 6B), demonst
rating the applicability of object detection networks for mode-of-action-based drug screening.

### Denoising

High contrast and fast image acquisition are critical to capture the dynamic nature of biology in full detail. How-ever, these usually come associated with high power imaging regimes often not compatible with live-cell imaging. Several denoising techniques such as PureDenoise (46) or DL-based approaches, both self-supervised (e.g. Noise2Void (16)) and supervised (e.g. CARE (15)), have been proposed to circumvent this experimental paradox. As these approaches allow for faster and more gentle imaging, we consider them as powerful tools for bacteriology. To test the applicability of denoising approaches to bacterial data, we recorded paired low and high signal-to-noise ratio (SNR) images of an H-NS-mScarlet-I (47) fusion protein in live *E. coli* cells. H-NS decorates the bacterial nucleoid homogeneously under nutrient-rich growth conditions and maintains nucleoid association after chemical fixation (48). This allows the study of chromosome organisation and dynamics, an important field of bacterial cell biology. We trained the CARE and N2V models on image pairs acquired using chemically fixed cells to prevent motion blur in the training dataset (Supplementary Table 9). We found that both parametric and DL-based approaches strongly increased the SNR of noise-corrupted images, as indicated by the peak signal-to-noise ratio (PSNR) and structural similarity index (SSIM) (49) (Figure 4A, right panel). These metrics are commonly used to access SNR and quality of image pairs, with higher values representing improved performance (Supplementary Note 1).

**Fig. 4.**
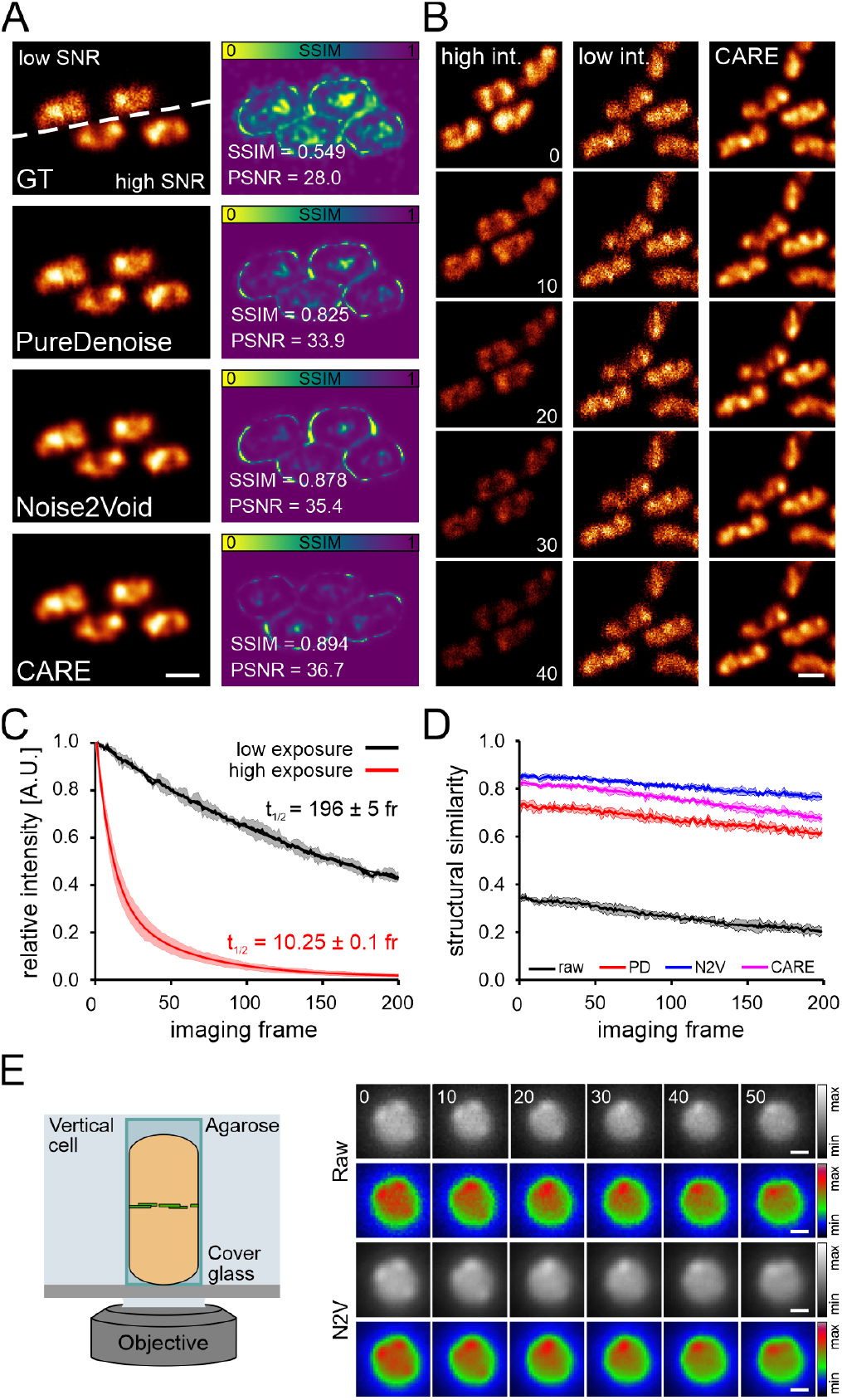
Image denoising for improved live-cell imaging in bacteriology. **[A]** Low and high signal-to-noise ratio (SNR) image pairs (ground truth, GT) of fixed *E. coli* cells, labeled for H-NS-mScarlet-I. Denoising was performed with PureDenoise (parametric approach), Noise2Void (self-supervised DL) and CARE (supervised DL). Structural similarity (SSIM) maps (right panel) comparing low-SNR or predictions to ground truth (GT) high-SNR data. SSIM and Peak-signal-to-noise ratio (PSNR) values are mean values of the two test images. **[B]** 10s interval representative time points of a live-cell measurement recorded at 1 Hz frame rate, demonstrating CARE can provide prolonged imaging at high SNR using low-intensity images as input. t_1/2_ represents the decay half time. A.U. = arbitrary units. **[C]** Intensity over time for different imaging conditions providing low/high SNR images shown in [A/B]. **[D]** Structural similarity between subsequent imaging frames (see Methods) was calculated for raw and restored time-lapse measurements. **[E]** Denoising of FtsZ-GFP dynamics in live *B. subtilis*. Cells were vertically trapped and imaged using the VerCINI method (41). Details are restored by Noise2Void (N2V), rainbow colour-coded images were added for better visualisation. [C] and [D] show mean values and respective standard deviations from 3 measurements. Scale bars are 1 µm.

Under the conditions tested, we obtained the best results using the supervised network CARE (SSIM = 0.894, PSNR = 36.7). We next applied the trained models to denoise live-cell time series recorded at low SNR conditions (Figure 4B, Supplementary Video 6). This led to an apparent increase in SNR, and intensity analysis revealed a 20x lower photobleaching rate as indicated by the exponential intensity decay time t_1/2_ (Figure 4C). However, the performance of the different denoising approaches in fast live cell measurements could not be assessed by the standard PSNR and SSIM metrics due to the lack of paired high-SNR images. As noise reduces contrast and structural information content in images, subsequent image frames in low-SRN time series should exhibit higher signal variation than in their high-SNR counterparts. We hence speculated that calculating the structural similarity between successive image frames (e.g. between frame 1 and frame 2, frame 2 and frame 3, etc.) could report on denoising performance for live-cell time series (Figure 4D). Indeed, all denoising approaches significantly increased SSIM values while preserving relative intensities over time (Supplementary Figure 7A).

As all the previous models were trained on fixed-cell data, the results on live-cell data could be compromised by potential fixation artefacts. Because N2V is self-supervised, it was possible to train it directly on the live-cell data. Hence, we could compare the performance of a N2V model trained on fixed cell images with the one trained on live-cell images. This resulted in a high structural similarity over the entire time series (Supplementary Figure 7B), indicating that no artefacts were introduced by training on fixed-cell data. Similar observations were made when comparing the fixed-cell N2V and CARE models. Analysis of raw and denoised fixed-cell time series using CARE further showed a constant SSIM value of 0.96 in the subsequent frame analysis (Supplementary Figure 7C). This indicates that the high contribution of shot-noise under low-SNR conditions can be overcome by the denoising method. Of note, the SSIM value obtained in fixed-cell measurements is higher than for denoised live-cell time series (0.82) (compare Figure 4D and Supplementary Figure 7C). To test whether this effect is caused by nucleoid dynamics (Supplementary Video 6), we recorded a time series under high-SNR imaging conditions using a small region of interest. High SNR leads to lower contribution of noise and higher SSIM values for subsequent image frames. As expected, the high laser power and longer exposure time induced strong photobleaching, leading to a rapid drop in structural similarity over time (Figure 4B, Supplementary Figure 7D). However, the first SSIM value of the high-SNR time series (representing the similarity between frame 1 and frame 2) is close to the corresponding SSIM value of the denoised low-SNR time series. This indicates that (i) the model provides optimal denoising performance and that (ii) the lower SSIM values in live-cell measurements originate from nucleoid dynamics rather than representing denoising artefacts.

As another example, we denoised time-lapse images of FtsZ treadmilling in live *B. subtilis* cells. These movies were recorded in vertically aligned cells using the so-called VerCINI approach (Figure 4E) and contributed to study the critical role of FtsZ treadmilling in cell division (41). As the constant movement of FtsZ-GFP renders acquisition of low and high SNR image pairs difficult, we used the self-supervised N2V method for the denoising task. Here, denoising emphasises subtle details that are difficult to be identified in the raw image data (Figure 4E). This allows for long time-lapse imaging of FtsZ dynamics with enhanced image quality (Supplementary Video 7).

### Artificial labelling and super-resolution prediction

Artificial labelling creates pseudo-fluorescent images from bright field, histology or EM images (13, 14, 50). It is beneficial for bright field-to-fluorescence transformation, as it does not require fluorophore excitation, being even less phototoxic than denoising of low SNR images while providing molecular specificity. Here, the neural network learns features in bright field images imprinted by specific structures or biomolecules (for example, membranes or nucleic acids) and creates a virtual fluorescence image of these structures. In contrast to image segmentation, artificial labelling does not require manual annotation, which reduces the time required for data curation. In the original published work, this allowed predicting multiple subcellular structures from single bright field stacks in mammalian tissue culture samples (13, 14). Because of a much smaller size, bacteria provide less structural information in bright field images than eukaryotic cells (Figure 5A). However, we regard the cell envelope as a promising target for artificial labelling, as exact cell shape determination is of interest for morphometric studies (e.g. for antibiotic treatments) or investigating the precise positioning of target molecules in individual cells (51–53). This is even more valuable if super-resolution information is obtained. To explore whether such information can be extracted using DL, we recorded different training datasets. The first dataset includes bright field and corresponding diffraction-limited widefield fluorescence images, in which the *E. coli* membrane is stained by the lipophilic dye Nile Red (Figure 5A, upper row). For the second dataset, we acquired super-resolved PAINT images (54, 55) and upsampled the respective brightfield image to match the resolution of the super-resolved membrane images (Figure 5A, lower row). For both datasets, we tested a 2D version of fnet (14), as well as CARE. For the diffraction-limited dataset, both networks were able to predict pseudo-fluorescence images from bright field images, with fnet showing slightly better performance according to the structural similarity metric (SSIM_fnet_ = 0.88 ± 0.06, SSIM_CARE_ = 0.83 ± 0.05) (Figure 5A). This is not surprising as fnet was designed for artificial labelling, while the good performance of CARE demonstrates the versatility of this network. Similar values were obtained for the super-resolution dataset (Figure 5A, Supplementary Figure 8), with predictions showing good agreement also on the sub-diffraction level (see cross-section as inset in Figure 5A). Additionally, although trained on fixed cells, the model can also be used to predict highly resolved membrane signal in live-cell time series (Supplementary Video 8). We then wanted to know how well our model generalises, i.e. whether it can predict the super-resolved membrane of bacteria grown in the presence of different antibiotics (see methods). Both fnet and CARE models successfully predicted the membrane stains in drug-treated cells (SSIM 0.8 - 0.9), indicating that it detects image features independent from the cell shape (Supplementary Figure 8). The increased resolution and contrast in membrane predictions allow to map the positioning of subcellular structures (here, the nucleoid) with higher precision (Figure 5B). This typically requires the acquisition of multi-colour super-resolution images (55), which is far more intricate and time-consuming.

**Fig. 5.**
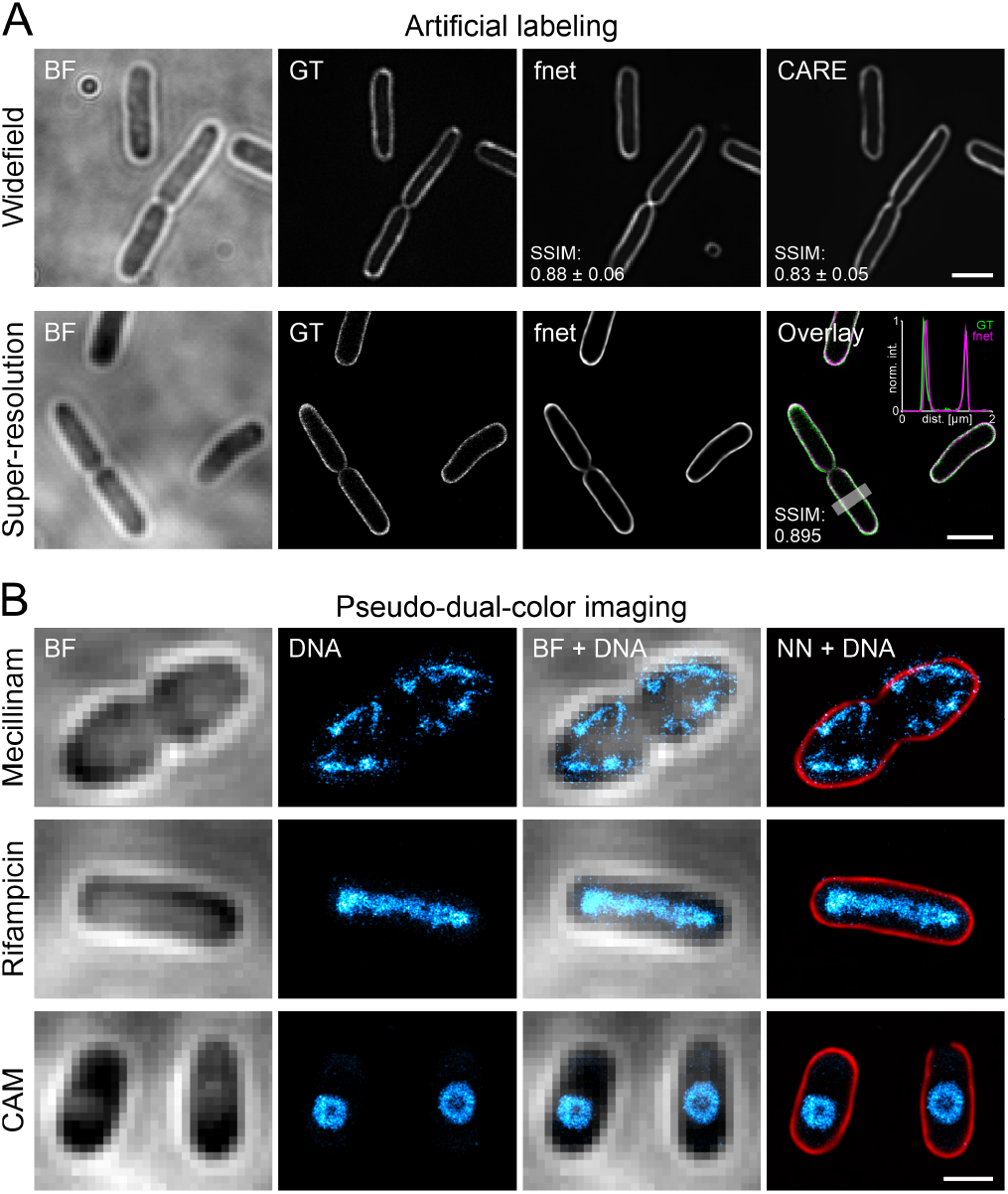
Artificial labelling of *E. coli* membranes. **[A]** fnet and CARE predictions of diffraction-limited (top row) and PAINT super-resolution (bottom row) membrane labels obtained from bright field (BF) images. GT = ground truth. **[B]** Pseudo-dual-color images of drug-treated *E. coli* cells. Nucleoids were super-resolved using PAINT imaging with JF646-Hoechst. Membranes were predicted using the trained fnet model. CAM = Chloramphenicol. Scale bars are 2 µm (A) and 1 µm (B).

Super-resolution membrane images can also be obtained by training a supervised DL network on paired low-resolution/high-resolution image datasets (15, 17, 18, 23). Here, we used structured illumination microscopy (SIM) (56) to record membrane images of dye-labelled live *E. coli* and *S. aureus* cells. SIM images are reconstructed from a set of images recorded at different grid positions and angles, which hence requires higher light doses than a single widefield image. As acquisition of such image sets is only required during the network training, but not during its application, super-resolution prediction reduces the light dose and also increases the achievable temporal resolution. Training of two CARE models on paired low/high-resolution images of *E. coli* and *S. aureus* using the ZeroCostDL4Mic notebook provided models that facilitate robust prediction of SIM images from single widefield snapshots (Figure 6). Here, contrast and resolution of predictions agreed well with reconstructed SIM images (Supplementary Video 9), as shown for cross sections along single *E. coli* and *S. aureus* cells (Figure 6, bottom panel). To evaluate the results, we performed SQUIRREL analysis, which detects reconstruction artefacts in super-resolution images (57). This analysis yielded similar errors for neural network predictions compared to SIM reconstructions, both for *E. coli* (resolution-scaled Pearson coefficient of 0.898 ± 0.018 (SIM) vs 0.907 ± 0.018 (prediction)) and *S. aureus* (0.957 ± 0.012 (SIM) vs 0.963 ± 0.010 (prediction)) (Supplementary Figure 9). SSIM values between predictions and GT SIM images were determined as 0.84 ± 0.03 (*E. coli*) and 0.92 ± 0.01 (*S. aureus*). Estimating the spatial resolution using image decorrelation (58) verified the very good agreement between the predicted (137 ± 7 nm for *E. coli* and 134 ± 5 nm for *S. aureus* images) and reconstructed SIM images (122 ± 2 nm for *E. coli* and 134 ± 1 nm for *S. aureus* images) with the expected 2x increase in resolution (308 ± 24 nm for *E. coli* and 289 ± 5 nm for *S. aureus* widefield images). This strategy is hence well suited to perform single-image super-resolution microscopy (18) in bacterial cells.

**Fig. 6.**
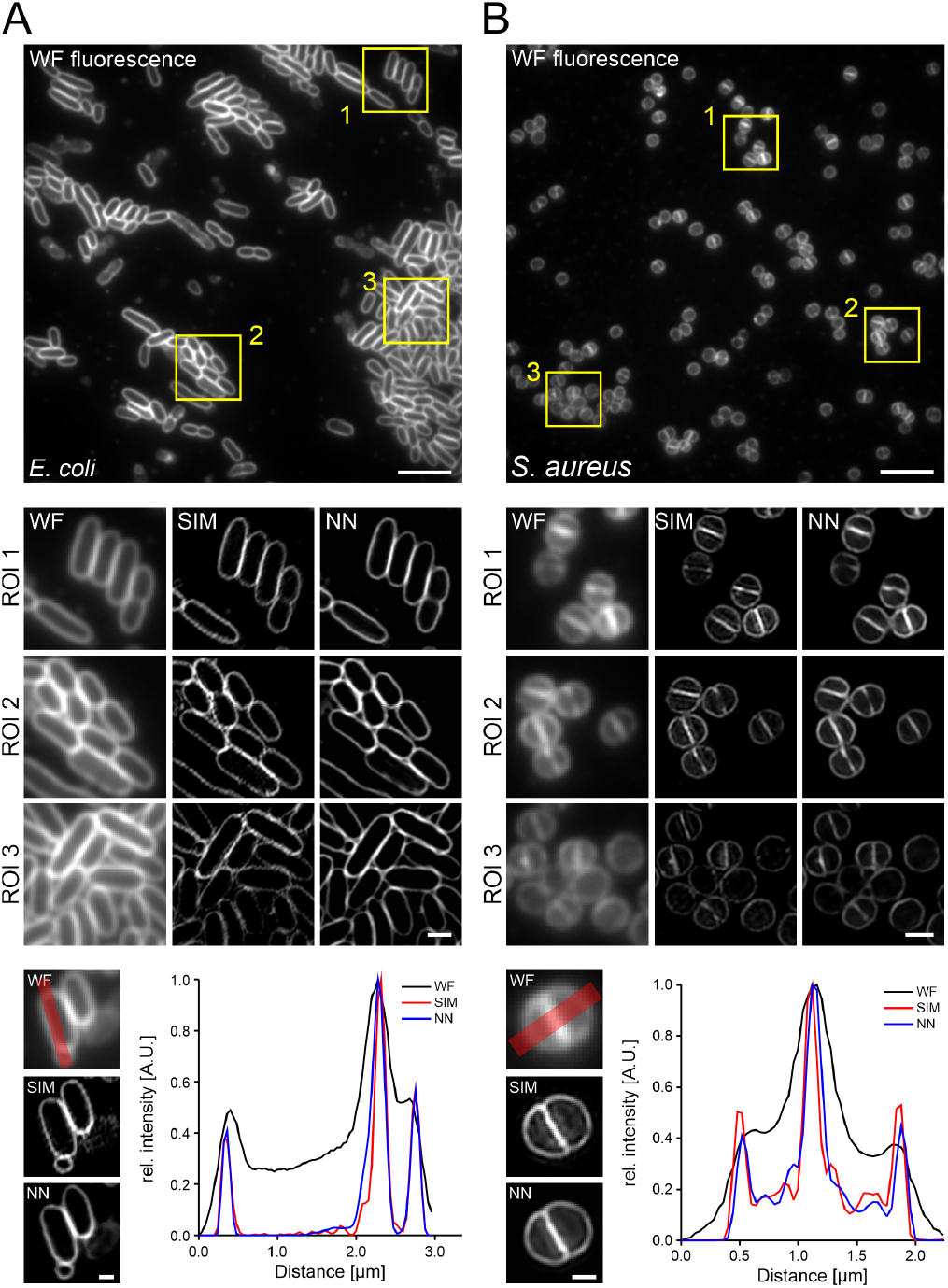
Prediction of SIM images from widefield fluorescence images. Widefield-to-SIM image transformation was performed with CARE for [A] live *E. coli* (FM5-95) and [B] *S. aureus* (Nile Red) cells. WF = widefield; NN = neural network output. Line profiles show a good agreement between prediction and ground-truth (bottom panel). Scale bars are 10 µm (overview images), 1 µm (magnified areas) and 0.5 µm (bottom panel)

## Discussion

In this work we demonstrate the potential of open-source DL technology for the analysis of bacterial bioimages. We employ popular DL networks that were developed by the open-source research community and are implemented in, but not limited to, the user-friendly ZeroCostDL4Mic platform (23). This enabled us to perform a variety of different image analysis tasks, such as image segmentation, object detection, image denoising, artificial labelling and the prediction of super-resolution images (Figure 1). Using the datasets that we provide, well-performing models can be trained within the time course of hours (see Supplementary Tables 3, 5, 9-11). Depending on the network, several tens of input images were sufficient, showing that valuable models can be generated even with a limited dataset size and thus moderate effort in data curation. For example, we annotated 7 images for segmentation of *S. aureus* cells (divided in 28 patches, in total 1249 cells), which required approximately 6 hours of manual work. Still, the trained StarDist model detected 98% of the cells in the test dataset of membrane-stained *S. aureus* (Figure 2). We think that this is an important point, as many researchers assume that huge training datasets are required for DL, often leading to discouragement and preventing the researcher from entering the field.

Another hurdle for new researchers is that the number of DL networks for a specific task continuously increases, and their performance can vary strongly depending on the images to be analysed (59). User-friendly implementations allow researchers to test the different networks and identify the best-performing network for a particular dataset. Testing different segmentation networks, we found StarDist and SplineDist being well suited to segment small rod-shaped and coccoid bacteria in bright field and fluorescence images (Figure 2, Supplementary Figure 3), while U-Net and pix2pix performed better for elongated cells at low to mid cell density. The performance at higher densities could be improved by predicting cell boundaries and cytosol using a multilabel U-Net, followed by post-processing of the network output. Thus, having a closer look at the input data can already give indications about which network might be more or less suited for the segmentation task (Supplementary Note 2). In our experience, the networks explored in this work are well suited to segment images recorded under standard conditions (e.g. exponential growth phase, regularly shaped cells, narrow size distribution). However, they might be of limited use or require large training datasets for more specialised cases, e.g. studying filamentation, irregularly shaped cells or biofilms. In such cases, we refer to DL networks developed for the particular segmentation task (29, 30, 32, 34, 35). Instance segmentations can subsequently be used for downstream applications such as tracking cell lineages or morphological changes. If this is not already included in the network (31, 35), segmentation masks can be used with the Fiji plugin TrackMate, which was recently updated for the use of DL technology (40) (Supplementary Video 3).

For object detection, we successfully trained models to detect and discriminate cells in specific growth stages (Figure 3A) or treated with different antibiotics (Figure 3B). Next to their use in image analysis, trained models can also be integrated into smart imaging pipelines in which the microscope system autonomously decides when and/or how to image a particular region of interest. Triggers can be the presence (or dominance) of a specific class (43) or the occurrence of class transitions (for example initiation of cell division). We anticipate this to be particularly powerful to study rare events, as smart acquisition strongly reduces data waste and data curation time (43, 60). At the same time, AI-based antibiotic profiling bears great promises for drug-screening applications and antibiotic mode of action studies. Although being trained on a limited dataset, the YOLOv2 model was able to discriminate between different antibiotic treatments based on their phenological fingerprint. We demonstrated that it could already be used for drug screening applications, as it was able to predict a similar mode of action (rod-to-sphere transition) for MP265-treated cells when trained on Mecillinam-treated cells (Supplementary Figure 6B). As training an object detection network only requires drawing of bounding boxes and no intensive feature design, it can be used straightforwardly by researchers new to this field, especially as membrane and DNA stains are widespread and easy to use. However, we think that the predictive power can be further improved by adding more fluorescent channels, such as indicator proteins that for example report on membrane integrity or the energetic state of the cell. This will result in comprehensive models that can be employed for automatic screening of large compound libraries (3) and might contribute to the discovery of novel antimicrobial compounds, which is desperately needed to tackle the emerging antibiotic resistance crisis (61). For cell-biological studies in microbiology, denoising represents a universally applicable strategy (62). Supervised DL networks are preferred over self-supervised ones (Figure 4), if well-registered image pairs can be acquired (Figure 4). This is mostly the case for static or slow-moving targets, but acquisition of training data on fixed specimens represents a good alternative (15). We note, however, that this requires proper controls to exclude fixation artefacts (as we have done in previous work (55, 63) (Supplementary Figure 7C)). Artefacts might be learned by the model and erroneously introduced into live-cell data during prediction, which can be hard to detect and lead to misinterpretation. Overall, all denoising approaches strongly improved the quality and SNR of nucleoid images in fixed cells (Figure 4A), with CARE (supervised DL) expectedly outperforming N2V (self-supervised) and PureDenoise (parametric) on our test dataset. Using the trained CARE model on labelled *E. coli* nucleoids in fast-growing cells revealed both high nucleoid complexity and dynamics on the second time-scale (Supplementary Video 6). We observed high-density regions which dynamically move within the area populated by the nucleoid. Such ‘super-domains’ were reported in previous studies (64, 65), eventually representing macrodomains or regions of orchestrated gene expression. Strong chromosome fluctuations were also reported in cells with a long generation time, showing an elongated nucleoid positioned along the bacterial long axis (66). In contrast, a highly complex nucleoid with structural stability on the minute time scale was observed during fast growth (67). Our data suggests that fast dynamics and a stable global nucleoid structure likely coexist. This might be important to maintain nucleoid flexibility while at the same time allowing for robust chromosome segregation during fast growth. The molecular or physico-chemical basis for this, however, needs to be further investigated.

When acquisition of high-quality data is challenging and no paired high-SNR images are available, self-supervised networks such as Noise2Void can be employed (62). We show this for time-series of FtsZ-GFP in vertically aligned *B. subtilis* cells (41), in which the gain in SNR allows following subtle FtsZ structures during their treadmilling along the cell septum (Figure 4E, Supplementary Video 7). Thus, even without access to high-SNR ground truth data, denoising can substantially increase image quality in challenging live-cell data. Concluding, we see the largest benefit of denoising in long-term microscopy experiments and capturing of fast dynamics. These experiments are strongly limited by phototoxic effects, photobleaching and in temporal resolution, parameters that are improved by denoising approaches.

We further consider artificial labelling and prediction of super-resolution images as future approaches with broad applicability. Increasing specificity and spatial resolution of bioimages (13, 14, 17, 18) is particularly useful in small bacterial cells, where most processes occur on scales close to or below the diffraction limit. Training a CARE model on paired bright field and super-resolution membrane images, we were able to artificially label membranes with subpixel accuracy (Figure 5A). Precise (sub-diffraction) prediction of cell boundaries enables the determination of cell size and shapes (morphometric analysis), which is important to describe and compare deletion mutants, drug-treated cells or cells grown under different environmental conditions (68). In subcellular localisation studies, fluorescence images are often overlaid with the respective bright field image. This provides only a very inaccurate picture of the relative molecule localisation, which can be strongly improved using our artificial labelling approach (Figure 5B). Additionally, as artificial labelling does not require a fluorescent label, it further opens up a spectral window for other fluorescent targets, thus increasing multiplexing capabilities. Using membrane stains, DL can be efficiently used to increase the spatial resolution, as we showed by predicting SIM images from diffraction-limited fluorescence signal using supervised learning. The enhanced resolution can improve downstream applications such as analysis of cell cycle stages in spherical bacteria (69). As bright field or fluorescence membrane images are part of basically any study including microscopy data, we think that artificial labelling and prediction of super-resolution images can be very useful for the bacterial research community.

As a general but important note, DL models are highly specific for the type of data they were trained on. Evaluating the model on ground truth data is thus essential to validate model performance, identify potential artefacts and and avoid a replication crisis for DL-based image analysis (59). Already slightly varying image acquisition parameters can transform a model from a good performer to a source of artefacts. Such parameters include different magnifications (pixel sizes), varying focal planes, illumination patterns, camera settings, and many more. However, even if pretrained models do not provide satisfying results, they can be used for transfer-learning, which can strongly accelerate the training and increase model performance. Collecting pretrained models in model zoos (such as the BioImage model zoo: https://bioimage.io/ can create a database encompassing a variety of species, microscopy techniques and experiments. This database can be used by the researchers to explore potential DL applications and apply pretrained models to their own research using designated platforms (9, 15, 23, 25, 38). Together with easily accessible DL networks and shared datasets, this work can support researchers to familiarise themselves with DL and find an entry point into the DL universe. Datasets and models generated in this work can be downloaded via Zenodo (see Supplementary Table 2), while further documentation on sample preparation, data preprocessing, training parameters and example images can be found on our GitHub repository (https://github.com/HenriquesLab/DeepBacs/wiki). In summary, user-friendly DL approaches possess large potential for image analysis in bacteriology. Both alone and in combination with classical image analysis routines, DL approaches will help to increase the amount and quality of information that can be extracted from bacterial bioimages.

## Availability

Datasets and models are available on Zenodo (Supplementary Table 2). Notebooks can be accessed via the ZeroCostDL4Mic repository (https://github.com/HenriquesLab/ZeroCostDL4Mic/wiki).

## ACKNOWLEDGEMENTS

C.S. and M.H. acknowledge funding by the Deutsche Forschungsgemeinschaft (German Science Foundation; DFG), grants HE 6166/17-1 and SFB 1177. C.S. further acknowledges support by the European Molecular Biology Organization (EMBO) in form of a Scientific Exchange Grant (grant nr. 8587). R.F.L. would like to acknowledge the support of the MRC Skills development fellowship (MR/T027924/1). P.M.P acknowledges funding by a La Caixa Junior Leader Fellowship (LCF/BQ/PI20/11760012) financed by “la Caixa” Foundation (ID 100010434) and by European Union’s Horizon 2020 research and innovation programme under the Marie Sklodowska-Curie grant agreement No 847648. Further, P.M.P. acknowledges funding by a Science and Technology Foundation grant (PTDC/BIA-MIC/2422/2020) and by Project LISBOA-01-0145-FEDER-007660 Microbiologia Molecular, Estrutural e Celular (to ITQB-NOVA). M.G.P. acknowledges funding by the European Research Council (ERC-2017-CoG-771709). SH acknowledges funding support by a Wellcome Trus & Royal Society Sir Henry Dale Fellowship (206670/Z/17/Z). MC supported by a UK Medical Research Council doctoral studentship. G.J. was supported by grants awarded by the Finnish Cancer Organization, the Sigrid Juselius Foundation, the Academy of Finland (338537), the Åbo Akademi University Research Foundation (CoE CellMech; to G.J.), and the Drug Discovery and Diagnostics strategic funding to Åbo Akademi University. E.G.M. and R.H. are supported by Gulbenkian Foundation and received funding from the European Research Council (ERC) under the European Union’s Horizon 2020 research and innovation programme (grant agreement No. 101001332) (to R.H.), the European Molecular Biology Organization (EMBO) Installation Grant (EMBO-2020-IG-4734)(R.H.) and the Wellcome Trust (203276/Z/16/Z) (R.H.). The authors thank Alexandre Bisson for sharing *Agrobacterium tumefaciens* live-cell data and Kevin D. Whitley for providing *B. subtilis* FtsZ-GFP data.

## AUTHOR CONTRIBUTIONS

C.S, M.H. and R.H. conceived the project; L.v.C., R.F.L., E.G.S., G.J., and R.H. wrote source code in the ZeroCostDL4Mic project; C.S., P.M.P. and M.C. performed the image acquisition of the training and test data; C.S. and M.C. annotated the data. E.G.S. helped with model training and data analysis. C.S. wrote the manuscript with input from all co-authors.

## Methods

### Segmentation of *E. coli* bright field images

*E. coli* MG1655 cultures were grown in LB Miller at 37°and 220 rounds per minute (rpm) overnight. Working cultures were inoculated 1:200 and grown at 23°and 220 rpm to OD600 0.5 – 0.8. For time lapse imaging, cells were immobilised under agarose pads prepared using microarray slides (VWR, catalogue number 732-4826) as described in de Jong et al., 2011 (1). Bright field time series (1 frame/min, 80 min total length) of 10 regions of interest were recorded with an Andor iXon Ultra 897 EMCCD camera (Oxford instruments) attached to a Nikon Eclipse Ti inverted microscope (Nikon Instruments) bearing a motorized XY-stage (Märzhäuser) and an APO TIRF 1.49NA 100x oil objective (Nikon Instruments). To generate the segmentation training data, 19 individual frames from different regions of interest were rescaled using Fiji (2x scaling without interpolation) to allow for betterer annotation. Resulting images were annotated manually using the freehand selection ROI tool in Fiji. For quality control, a test dataset of 15 frames was generated similarly. Contrast was enhanced in Fiji and images were either converted into 8-bit TIFF (CARE, U-Net, StarDist) or PNG format (pix2pix).

### Data pre- and post-processing for cell segmentation using the multi-label U-Net notebook

In order to improve segmentation performance, we employed a U-Net that is trained on semantic segmentations of both cell cytosol and boundaries. To generate the respective training data, annotated cells were filled with a gray value of 1, while cell boundaries were drawn with a gray value of 2 and a line thickness of 1. Together with the fluorescence image, this image was used as network input during training. During post-processing, cell boundaries were subtracted from predicted cell segmentations, followed by thresholding and marker-based watershed segmentation (Fiji plugin “MorpholibJ” (2)). Pre- and post-processing routines are provided as Fiji macros and can be downloaded from the DeepBacs github repository.

### Segmentation of *S. aureus* bright field and fluorescence images

For *S. aureus* time-lapse experiments overnight cultures of S. aureus strain JE2 were back-diluted 1:500 in TSB and grown to mid-exponential phase (OD_600nm_ = 0.5). One milliliter of the culture was incubated for 5 min (at 37°C) with the membrane dye Nile Red (5 µg/ml, Invitrogen), washed once with phosphate buffered saline (PBS), subsequently pelleted and resuspended in 20 µl PBS. One microliter of the labelled culture was then placed on a microscope slide covered with a thin layer of agarose (1.2% (w/v) in 1:1 PBS/TSB solution). Time-lapse images were acquired every 25 s (for DIC) and 5 min (for fluorescence images) by structured illumination microscopy (SIM) or classical diffraction limited widefield microscopy in a GE HealthCare Deltavision OMX system (with temperature and humidity control, 37°). The images were acquired using 2 PCO Edge 5.5 sCMOS cameras (one for DIC, one for fluorescence), an Olympus 60x 1.42NA Oil immersion objective (oil refractive index 1.522), Cy3 fluorescence filter sets (for the 561 nm laser) and DIC optics. Each time-point is a Z-stack of 3 epifluorescence images using either the 3D-SIM optical path (for SIM images) or classical widefield optical path (for non super-resolution images). These stacks were acquired with a Z step of 125 nm in order to use the 3D-SIM-reconstruction modality (for the SIM images) of Applied Precision’s softWorx software (AcquireSRsoftWoRx), as this provides higher quality reconstructions. A 561 nm laser (100 mW) was used at 11–18 W cm-2 with exposure times of 10-30 ms. For single-acquisition S. aureus experiments, sample preparation and image acquisition was performed as mentioned above but single images were acquired. To generate the training dataset for StarDist segmentation, individual channels were separated and preprocessed using Fiji (3, 4). Nile Red fluorescence images were manually annotated using ellipsoid selections to approximate the *S. aureus* cell shape. Resulting ROIs were used to generate the required ROI map images (using the “ROI map” command included in the Fiji plugin LOCI) in which each individual cell is represented by an area with a unique integer value. Training images (512 × 512 px^2^) were further split into 256 × 256 px^2^ images, resulting in 28 training images pairs. 5 full field-of-view test image pairs were provided for model quality control. For segmentation of S. aureus bright field images (corresponding to the Nile Red fluorescence images) were paired with the ROI masks created for fluorescence image segmentation.

### Segmentation of live *B. subtilis* cells

*Bacillus subtilis* cells expressing FtsZ-GFP were prepared as described in (Whitley *et al*., 2021) (5) (strain SH130, PY79 L)hag ftsZ::ftsZ-gfp-cam). Strains were taken from glycerol stocks kept at -80°and streaked onto nutrient agar (NA) plates containing 5µg/ml chloramphenicol then grown overnight at 37°C. Liquid cultures were started by inoculating time-lapse medium (TLM) (de Jong *et al*., 2011) (1) with a single colony and growing overnight at 30°with 200 rpm agitation. The following morning, cultures were diluted into chemically defined medium (CDM) containing 5µg/ml chloramphenicol to OD_600_ = 0.1, and grown at 30°C until the required optical density was achieved (5). All imaging was done on a custom-built, 100X inverted microscope. A 100x TIRF objective (Nikon CFI Apochromat TIRF 100XC Oil), a 200mm tube lens (Thorlabs TTL200) and Prime BSI sCMOS camera (Teledyne Photometrics) were used achieving an imaging pixel size of 65nm/pixel. Cells were illuminated with a 488 nm laser (Obis) and imaged using a custom ring-TIRF module operated in ring-HiLO (6). A pair of galvanometer mirrors (Thorlabs) spinning at 200Hz provides uniform, high SNR illumination (5). The raw data analyzed here were acquired and analysis of that raw data presented in Whitley et al., 2021 (5). These data have now been reanalyzed using cell segmentation methods discussed. Slides were prepared as described previously. Molten 2% agarose made with CDM was poured into gene frames (Thermo Scientific) to form flat agarose pads, then cut down to thin 5mm strips. 0.5 µl of cell culture grown to mid-exponential phase (OD_600_ = 0.2-0.3) was spotted onto the agarose and allowed to absorb (approx. 30 seconds). A plasma-cleaned coverslip was then placed atop the gene frame and sealed in place. Before imaging, the prepared slides were then prewarmed inside the microscope body at least 15 minutes before imaging. Time-lapse images were then taken in TIRF using a custom built 100x inverted microscope. Images were taken at 1 second exposure, 1 frame/minute at 1-8 W/cm2 (5). Videos were denoised using ImageJ plugin PureDenoise (7) then lateral drift was corrected using StackReg (8). To create the training dataset, 10 frames were extracted from each time-lapse approximately 10 frames apart. This was to ensure sufficient difference between the images used for training. Ground truth segmentation maps were generated by manual annotation of cells in each frame using the Fiji/ImageJ LabKit plugin lab (https://github.com/juglab/imglib2-labkit. This process assigns a distinct integer to all pixels within a cell region, and background pixels are labelled 0. A total of 4,672 cells were labelled across 80 distinct frames to create the final training dataset.

### Confocal imaging for denoising of *E. coli* time series

*E. coli* strain CS01 carrying a chromosomal H-NS-mScarlet-I protein fusion (parental strain NO34) was grown in LB Lennox at 25°and shaking at 220 rpm. To generate the training dataset, cells were fixed chemically using a mixture of 2% formaldehyde and 0.1% glutaraldehyde. Fixed or live cells were immobilized under agarose pads poured into gene frames following the protocol by de Jong et al. (1). Imaging was performed on a commercial Leica SP8 confocal microscopy (Leica Microsystems) bearing a 1.40 NA 63x oil immersion objective (Leica Microsystems). To increase optical sectioning, the pinhole size was set to 0.5 AU and 512 × 512 px^2^ confocal images (45 nm pixel size) were recorded and emission was detected with HyD detectors in standard operation mode (gain 100, detection window 570 – 650 nm). For the training dataset, a two-channel image of the same structure was recorded in frame sequential mode using different settings for low (0.03% 561 nm laser light, no averaging) and high SNR images (0.1% 561 nm laser light, 4x line averaging), respectively. For live-cell time series, the field of view was reduced to 256 × 256 px^2^ to allow for fast acquisition of high SNR images at 0.8 Hz. Low SNR time series were recorded at similar frame rate by including a lag time.

### Bacillus subtilis VerCINI microscopy

The raw data analysed here were acquired and analysis of that raw data is presented in Whitley et al. 2021 (5). These data have now been reanalysed using the denoising methods described. Silicone micropillar wafers were nanofabricated and used to prepare agarose microholes as described previously (5). Molten 6% agarose was poured onto the silicone micropillars and allowed to set, forming an agarose pad punctured with microscopic holes. The agarose pad was then transferred into a gene frame, and agarose surrounding the micro-hole array was cut away. Concentrated liquid cell culture at mid exponential phase (OD_600_ = 0.4) was loaded onto the pad and centrifugation using an Eppendorf 5810 centrifuge with MTP/Flex buckets loaded individual cells into the microholes. The pad was then washed to remove unloaded cells. This repeated several times until a sufficient level cell loading was achieved. Cells were imaged at 1 frame/second with continuous exposure for 2 minutes at 1-8 W/cm2 (5). Image denoising was performed using the Image J plugin PureDenoise (7) and lateral drift was then corrected using StackReg (8).

### *E. coli* cell cycle classification

Classification of rod-shaped, dividing and microcolonies was performed using the time series described in section **BL segmentation *E. coli***. Individual frames from several time series were used for training. To generate the training dataset, individual frames spread over the entire time series (typically frames 1, 15, 30, 55 and 80) were converted into PNG format. For the large field-of-view model, the entire image was used, while images were split into 4 regions of 256 × 256 px^2^ size for the small field-of-view model. Images were annotated using LabelImg (9). The final training dataset contained 28 annotated patches, and dataset size was increased 4x during training using data augmentation implemented in the ZeroCostDL4Mic YOLOv2 notebook (rotation and flipping).

### *E. coli* cell cycle classification

E. coli strain NO34 (10) was grown in LB at 32°shaking at 220 rpm overnight. Working cultures were inoculated 1:200 in fresh LB and grown to mid-exponential phase and antibiotics were added at the concentration and for the time listed in Supplementary Table 5. Antibiotic stock solutions were prepared freshly 5-10 min before use. Cells were fixed using a mixture of 2% formaldehyde and 0.1% glutaraldehyde, quenched using 0.1% sodium borohydrate (w/v) in PBS for 3 min and immobilized on PLL-coated chamber slides (see (11) for details). Nucleoids were stained using 300 nM DAPI for 15 min. After 3 washes with PBS, 100 nM Nile Red in PBS was added to the chambers and confocal images were recorded with a commercial LSM710 microscope (Zeiss, Germany) bearing a Plan-Apo 63x oil objective (1.4 NA) and using 405 nm (DAPI) and 543 nm (Nile Red) laser excitation in sequential mode. Images (800 × 800 px^2^) were recorded with a pixel size of 84 nm, 16-bit image depth, 16.2 µs pixel dwell time, 2x line averaging and 1 AU pinhole size.

4-8 confocal images were used to generate the training dataset, depending on the cell count per image (for example, only few cells are present per image for nalidixate treated cells, while many cells were present for chloramphenicol treatment). Each image was converted to PNG format, split into 4 non-overlapping patches (400 × 400 px^2^) and patches were annotated online using makesense.ai (Make Sense, n.d.). Annotations were exported in PASCAL VOC format. Next to the 5 antibiotic treatments and control conditions, vesicles and partially attached cells were added as additional classes (“Vesicles” and “Oblique”, respectively), resulting in a total of 8 classes. Synthetic test data was generated by randomly stitching 200 × 200 px^2^ patches of different drug treatments and the control condition. Small patches were manually cropped from images that were not seen by the network during the training. In total, 32 test images were generated this way and annotated online using makesense.ai as described above. Additionally, 400 × 400 px^2^ image patches of previously unseen images (drug-treatments and control) were annotated using LabelImg (9).

### Artificial labeling of *E. coli* membranes

PAINT super-resolution images of *E. coli* membranes were recorded as described elsewhere (11). In brief, cells were grown in LB at 37°C and 220 rpm, fixed in mid-exponential phase (OD_600_ = 0.5) using a mixture of 2% formaldehyde and 0.1% glutaraldehyde, immobilized on poly-L-Lysine coated chamber slides and permeabilized with 0.5% TX-100 in PBS for 30 min. 400 pM Nile Red in PBS was added and PAINT time series (6,000 - 10,000 frames) were recorded on a custom built setup for single-molecule detection (Nikon Ti-E body equipped with a 100x Plan Apo TIRF 1.49 NA oil objective) using 561 nm excitation ( 1 kW/cm^2^) or a commercial N-STORM system with a similar objective and imaging parameters. Two image datasets were recorded using either a 1x or 1.5x tube lens (158 and 106 nm pixel size, respectively). PAINT images were reconstructed using Picasso (13) and exported at different magnifications (8x for 158 nm pixel [19.8 nm/px] and 6x for 106 nm pixel size [17.7 nm/px]). Corresponding bright field images were scaled similarly in Fiji without interpolation and registered with the PAINT image. Multiple 512 × 512 px^2^ image patches were extracted from these images and used for model training. For artificial labeling in drug-treated cells, cells were exposed to the following antibiotics: 100 µg/ml rifampicin for 10 min, 50 µg/ml Chloramphenicol for 60 min, 2 µg/ml Mecillinam for 60 min. Further sample preparation and imaging was performed similar to untreated cells.

### Prediction of membrane SIM images in live *E. coli* cells

For widefield-to-SIM prediction experiments overnight cultures of *E. coli* strain DH5α were back-diluted 1:500 in LB and grown to mid-exponential phase (OD_600_ = 0.3). One milliliter of the culture was incubated for 10 min (at 37°) with the membrane dye FM5-95 (10 µg/ml, Invitrogen), washed once with PBS, subsequently pelleted and resuspended in 10 µl PBS. One microliter of the labelled culture was then placed on a microscope slide covered with a thin layer of agarose (1.2% (w/v) in 1:1 PBS/LB solution). Image acquisition was performed as mentioned in section **Segmentation of *S. aureus* bright field and fluorescence images**. To generate the paired training dataset for super-resolution prediction, raw SIM images were averaged to obtain the diffraction limited widefield image, while the in-focus plane of the SIM reconstruction was used as corresponding high-resolution image. The dataset was curated by removing defocused images and images with low signal resulting in reconstruction artefacts. In total, 55 training and 5 test image pairs were used.

### Calculation of the multiscale structural similarity index (SSIM)

Performance of several deep learning approaches (e.g. CARE) was accessed by calculating the multiscale structural similarity index (here denoted as SSIM) between the source/predicted image and the ground truth image (14) (see Supplementary Note 1). Since background is suppressed efficiently by most networks and is thus over-proportionally contributing to the average per-image SSIM value (leading to an over-optimistic value), we determined the SSIM only within the outlines of bacterial cells. For this, ROIs were generated in Fiji by thresholding the high SNR image or time series average image. For denoising of live-cell time series lacking ground truth data (e.g. N2V), we determined the SSIM value over time by comparing each image frame to the subsequent image frame of the time series (thus termed subsequent-frame SSIM). A low SSIM value thus depicts a high frame-to-frame variation.

### SQUIRREL analysis

To access artefacts in super-resolution prediction from widefield data we used the SQUIRREL algorithm implemented in the Fiji NanoJ plugin (15, 16). This way, the predictions of 5 WF images and the respective SIM ground truth images were analysed. SQUIRREL calculates a diffraction limited image from super-resolution images to compare them with the corresponding low-resolution ground truth image. Resulting error maps give rise to reconstruction and in this case also prediction artefacts.

