## Supplementary Information for "DeepBacs: Bacterial image analysis using open-source deep learning approaches"

### Supplementary Notes

#### Supplementary Note 1 - Prediction quality assessment metrics

Testing the performance of trained DL models is crucial to obtain reliable results. For this purpose, different metrics are commonly used, depending on the data type and DL task. All metrics used in this study are described in detail in the Supplementary Information of the ZeroCostDL4Mic publication (von Chamier *et al.*, 2021).

For segmentation tasks, we employed the Intersection-over-Union (IoU), Recall and Precision metrics.

*IoU*, also known as the Jaccard index, reports on the overlap between ground truth (GT) and predicted masks (P) and is calculated according to equation 1. A value of one thus represents perfect overlap, declining towards 0 with decreasing overlap.

$$IoU\ (GT, P)\ = \frac{|GT\ \cap\ P|}{|GT\ \cup\ P|} \qquad \text{(Eq. 1)}$$

While the IoU metric gives access to the performance of semantic segmentation (i.e. classification in the background and foreground pixels), it does not report on the quality of instance segmentation. A dataset might have a high IoU, but individual instances might not be resolvable (see CARE and pix2pix predictions of dense regions in Figure S1). Instead, we use the *Recall* and *Precision* metrics (Eq. 2 and 3, respectively), whose calculation is based on the true positives (TP), false negatives (FN), and false positives (FP) determined using the test dataset.

$$Recall\ = \frac{TP}{TP\ +\ FN} \qquad \text{(Eq. 2)}$$

$$Precision\ = \frac{TP}{TP\ +\ FP} \qquad \text{(Eq. 3)}$$

*Recall* thus describes the sensitivity of the analysis or prediction, while *Precision* describes the specificity.

In object detection, model performance is assessed using the Average Precision (AP). AP is calculated via so-called Precision-Recall-Curves. These curves are generated by applying different thresholds for object detection, followed by the calculation of Precision and Recall values for each threshold. Lowering the detection threshold leads to the detection of more instances (Recall) but typically leads to a reduced Precision, as also more false positives are detected. The AP value is finally received by determining the area under the curve. *mean Average Precision* (mAP) represents the mathematical mean of the AP values of all classes present in the test dataset.

The quality of image denoising was quantified using the structural similarity index (SSIM) (Wang *et al.*, 2004) and peak signal-to-noise ratio (PSNR). SSIM describes the similarity of two images A and B based on mean pixel intensities ($\mu_A$, $\mu_B$), pixel intensity variation ($\mu_A^2$, $\mu_B^2$) and pixel intensity covariance ($\sigma_{AB}$) (Equ. 4).

$$SSIM\,(A,B) \;=\; \frac{(2\mu_A\mu_B \;+\; C_1)(2\sigma_{AB} \;+\; C_2)}{(\mu_A^2 \;+\; \mu_B^2 \;+\; C_1)\,(\mu_A^2 \;+\; \mu_B^2 \;+\; C_2)} \qquad \text{(Eq. 4)}$$

Constants $C_1$ and $C_2$ were introduced for stability and are defined according to Equ. 5 and 6.

$$C_1 \;= (K_1L)^2 \qquad \text{(Eq. 5)}$$

$$C_2 \;= (K_2L)^2 \qquad \text{(Eq. 6)}$$

L represents the dynamic range of the images (maximal number of grey values), and $K_1$ and $K_2$ are constants that were set to 0.01 and 0.03 as suggested in the original publication (Wang *et al.*, 2004).

PSNR was originally introduced to compare the preservation of image quality in compressed images. However, PSNR can also be used to quantify denoising performance when ground-truth data is available. Its calculation is based on the mean squared error between two images, A (high SNR) and B (low SNR) (MSE, Eq. 7), and follows Eq. 8.

$$MSE \;=\; \frac{1}{n}\sum_{i,j}^{n}(A_{ij} \;-\; B_{i,j})^2 \qquad \text{(Eq. 7)}$$

with *n* representing the total number of pixels in the image and *i*, *j* the individual pixels.

$$PSNR \;= 20\,log_{10}\,(L) \;-\; 10\,log_{10}\,(MSE) \qquad \text{(Eq. 8)}$$

**Supplementary Note 2 - Selecting a network for bacterial segmentation**

Different DL networks were developed to perform semantic/instance segmentation of bioimages. The performance of these networks can vary strongly on the type of data (e.g. strains or microscopy technique) used.

*The following considerations can help to decide for a particular network:*

(i) Estimate the axial ratio of bacteria in the images. Small axial ratios (in our experience 1 -3; e.g. for slowly growing rod-shaped cells, stationary phase cells or cocci/ovococci) are ideal targets for StarDist independent of cell density.

(ii) Cells with axial ratios above ~4-5 or non-convex cells (e.g. fast-growing rod-shaped cells such as *E. coli* or *B. subtilis*) can be segmented with U-Net, CARE and pix2pix but require additional post-processing (thresholding and particle analysis). Examples of such processes are background subtraction, specific thresholding algorithms, pixel-based actions (e.g. dilation or erosion) active contour-based algorithms (Kass *et al.*, 1988), or marker-based watershed segmentation. The workflow for each model differs slightly (see Supplementary Figure 1), which also influences the time required for data preparation, up-/downloading and post-processing.

(iii) Prediction of multiple labels increases segmentation performance for densely growing cells. In this case, we suggest using the multi-label U-Net network that is trained on semantic images labelled for cell cytosol and boundaries.

(iv) To quickly get an impression of network suitability, we recommend transfer-learning using a few annotated images and a model pre-trained on images with similar parameters (pixel size, bacterial shape, image modality). Network performance should be evaluated using test data. Metrics such as IoU, Precision or Recall can be used for this purpose (see Supplementary Note 1) and are automatically calculated in the QC section of ZeroCostDL4Mic notebooks.

(v) There are many other segmentation approaches available that can be tested. For phase contrast images, for example, the MatLab-based SuperSegger (Stylianidou *et al.*, 2016) or Oufti (Paintdakhi *et al.*, 2016) can be employed. DeepLearning networks such as DeepCell (Van Valen *et al.*, 2016) or MISIC (Panigrahi *et al.*, 2021) can be used to segment and track cells at high density in phase contrast and for MISIC also in bright field images. Their implementation, however, might require more computational expertise or support from the developers.

#### Supplementary Figures

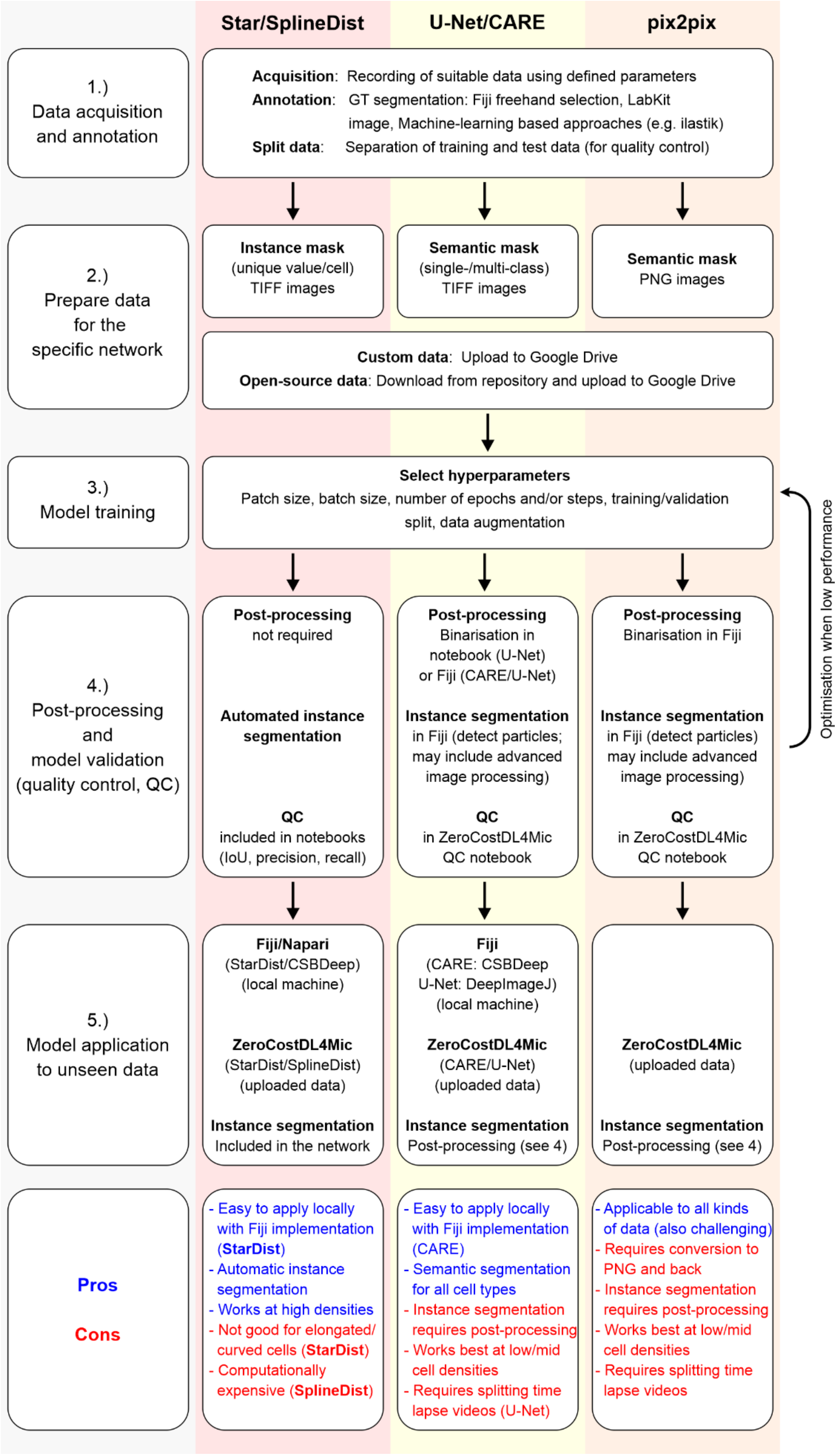


***Supplementary Figure 1****: Segmentation workflows using cloud implementations of different DL networks.*

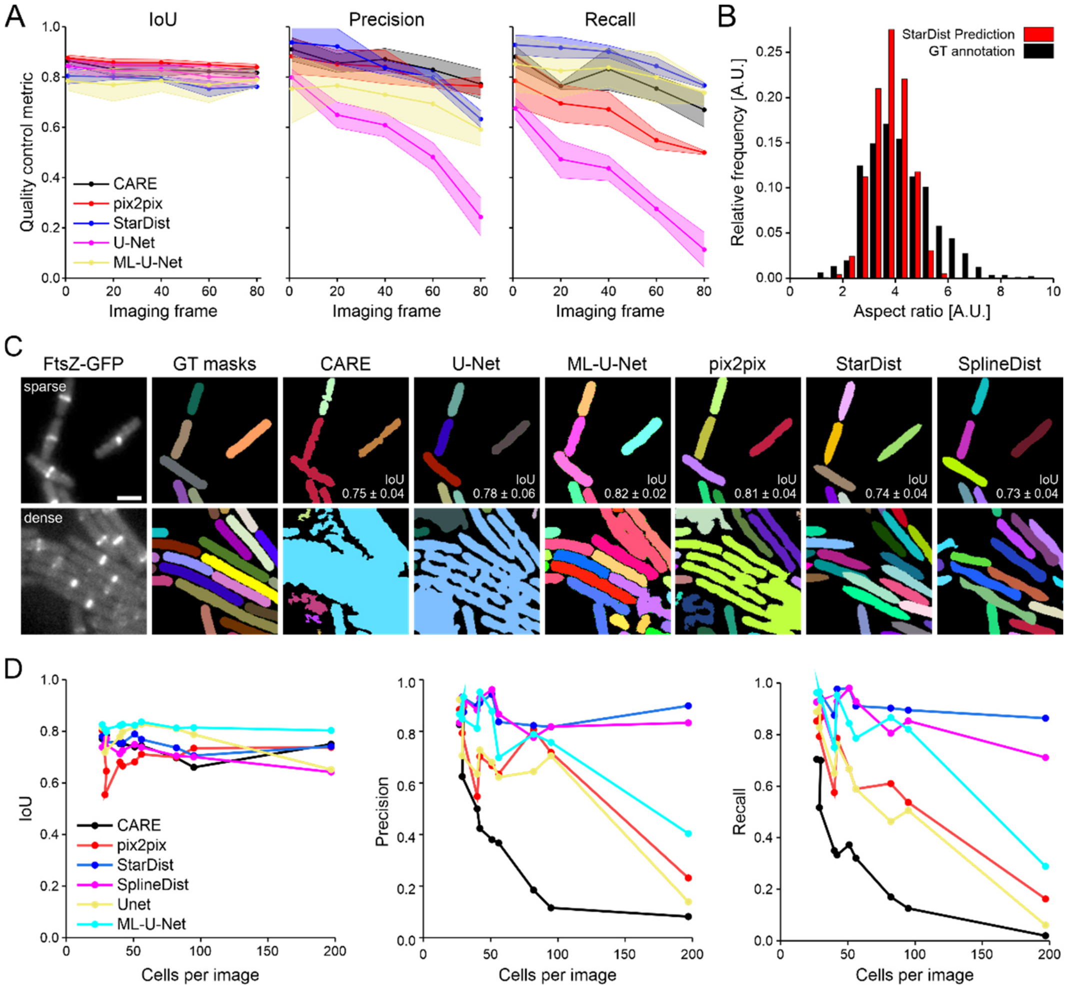


***Supplementary Figure 2**: Performance of brightfield and fluorescence image segmentation using different deep learning approaches. **[A]** Intersection over Union (IoU), precision and recall of instance segmentations of brightfield time-lapse images of growing E. coli cells generated with different networks. **[B]** Axial ratio analysis of StarDist instance segmentation of E. coli bright field images. **[C]** Segmentation of FtsZ-GFP fluorescence images of living B. subtilis. IoU values were determined for the test data set (10 image pairs). **[D]** Intersection over Union (IoU), precision and recall of instance segmentations of living, GFP-FtsZ expressing, B. subtilis cells generated with different networks. Scale bar in [C] is 2 µm*

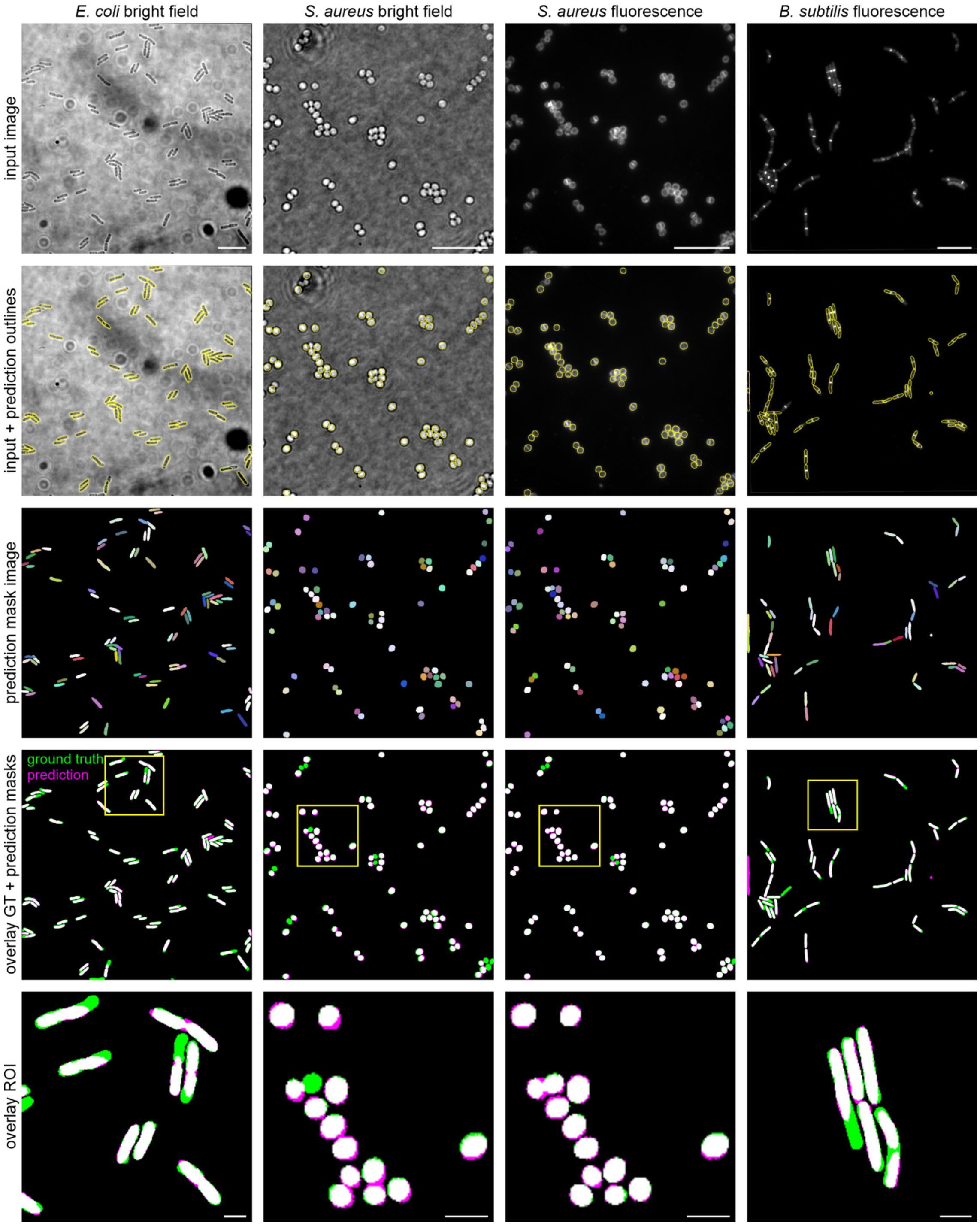


***Supplementary Figure 3***: *StarDist "master model" for segmentation of the four datasets provided in this work. The lowest panel shows insets of ground truth (GT) -prediction overlays marked by yellow rectangles in the overview image (second-last row). Scale bars are 10 µm (overviews) and 2 µm (ROIs).*

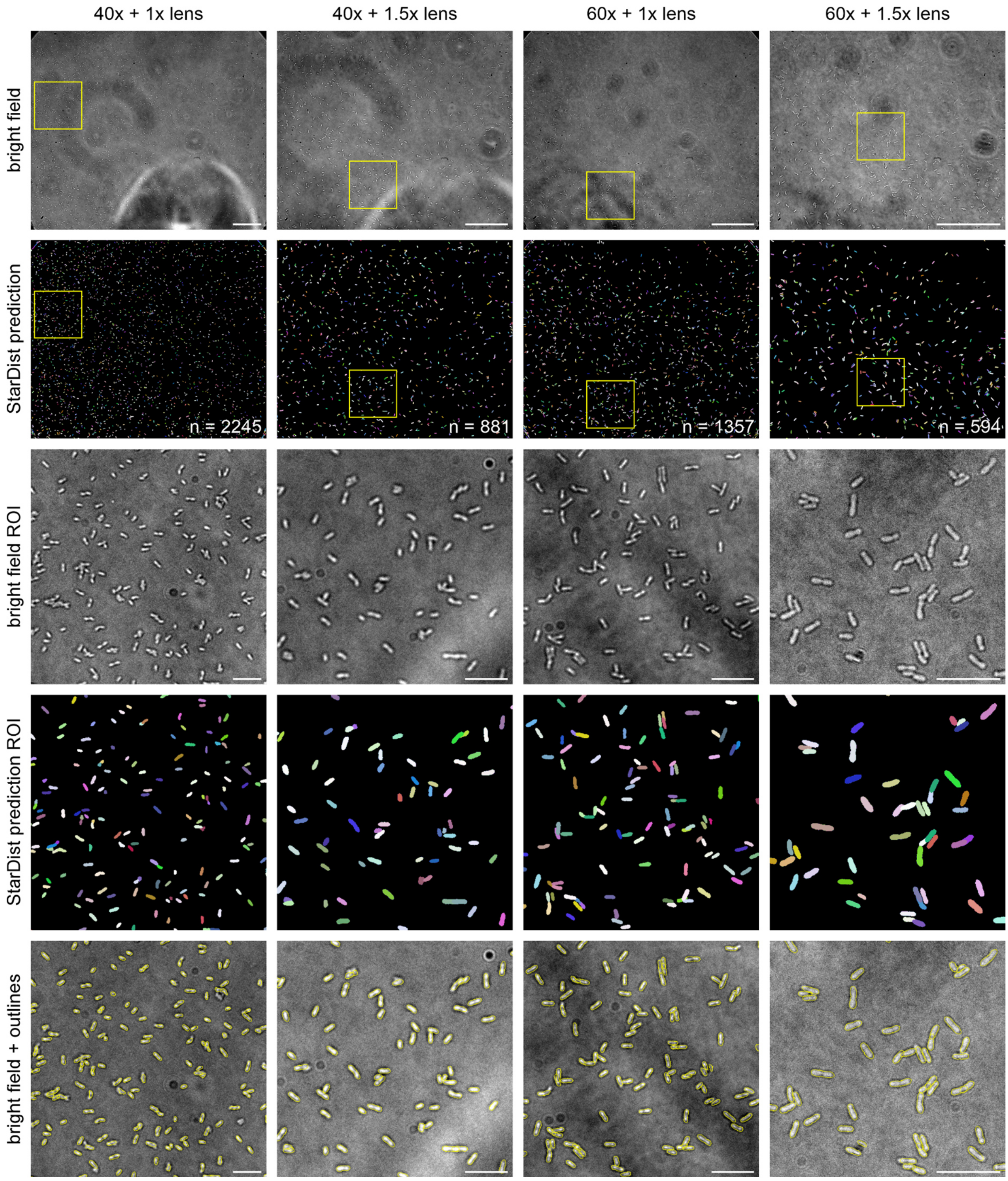


***Supplementary Figure 4****: Agrobacterium tumefaciens large field of view StarDist segmentations. A 2D StarDist model was trained on pooled bright field data (512 x 512 px patches) acquired with different objectives and at different magnifications. This allows predicting a high cell number in large images also at challenging bright field conditions. Scale bars are 20 μm (overview images) and 10 μm (ROIs).*

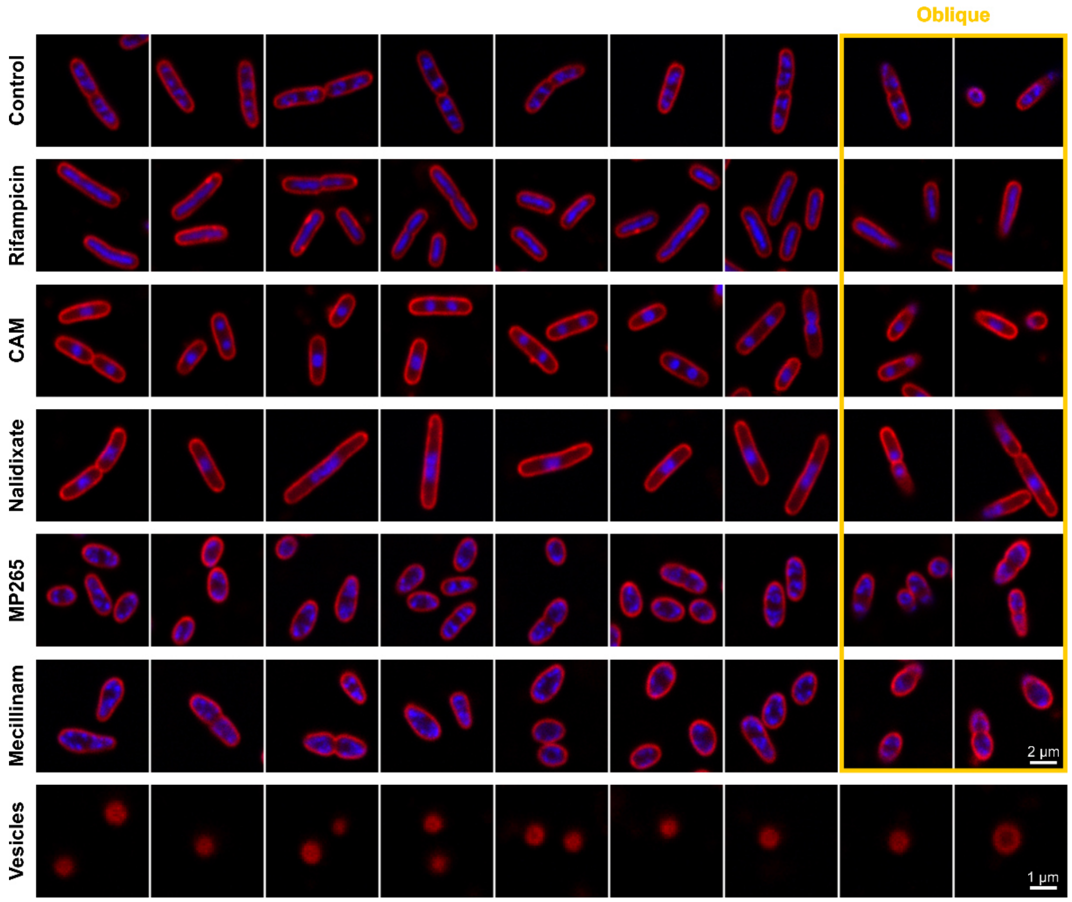


***Supplementary Figure 5****: Examples for the different classes used in DL-assisted antibiotic phenotyping. Oblique cells represent incompletely attached cells and are thus included within all antibiotic treatments and the control condition.*

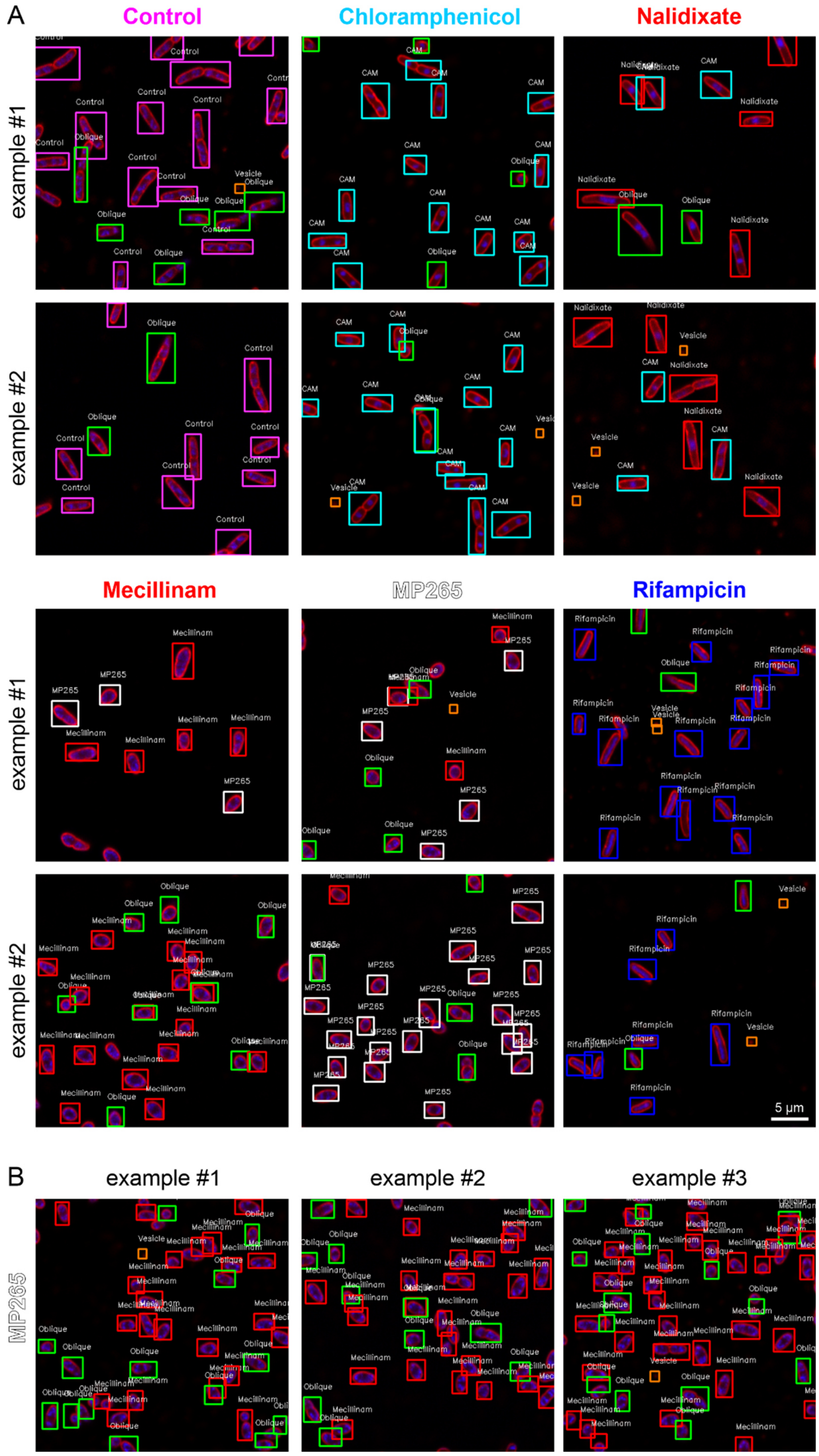


***Supplementary Figure 6****: Antibiotic profiling model predictions. [A] Representative regions of interest showing object detection results on the single-treatment test dataset. [B] Application of a YOLOv2 model to identify shared mode-of-actions. The model was trained on a dataset without MP265-treated cells. Application of the model to images with MP265-treated cells predicts Mecillinam treatment (red bounding box) due to a similar phenotype.*

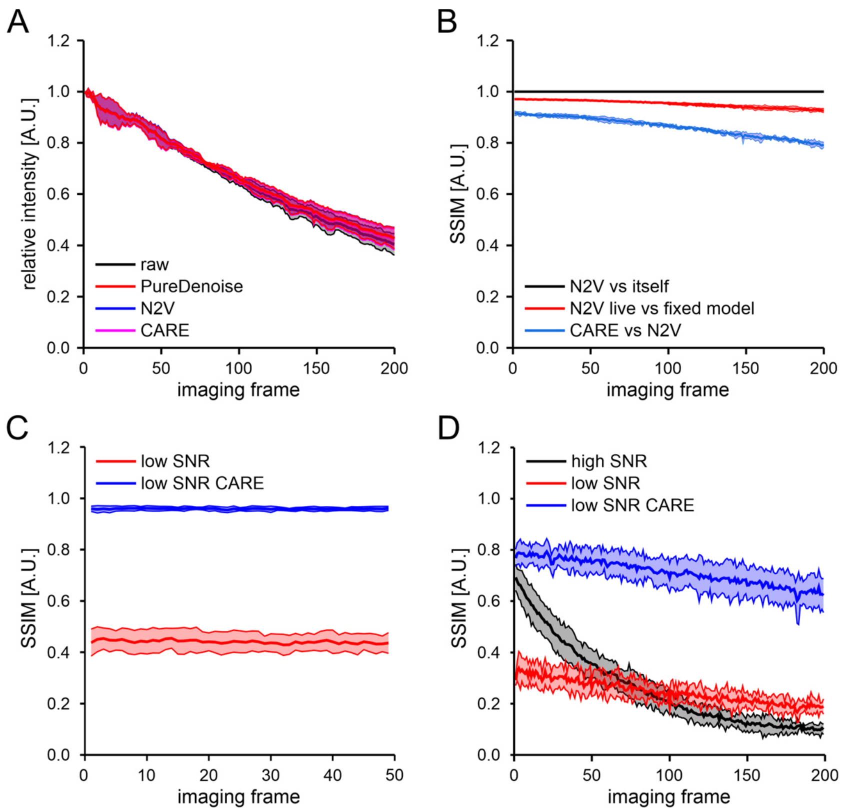


***Supplementary Figure 7**: Controls for DL-based image denoising. **[A]** Intensity over time for raw and denoised data. Denoising does not affect the relative intensities. **[B]** SSIM analysis of live-cell time series denoised with different approaches. Perfect similarity is given by a value of 1, as shown for SSIM analysis using the same time series (black line). A high similarity is observed between N2V models trained either on the fixed-cell dataset or on live-cell time series image frames. Time series denoised with CARE and N2V models also exhibit high SSIM values. Decreasing SSIM over imaging time probably arises from a better performance of supervised training (CARE) for images with lower SNR values, occurring at the end of the measurements due to photobleaching. **[C]** SSIM over time (subsequent image frames) of fixed cell time series, imaged at low SNR conditions (red) and denoised using CARE (blue). **[D]** SSIM over time (subsequent image frames) for high SNR (black, high laser power and 4x line averaging), low SNR (red, low laser power without averaging) and low SNR time series denoised with CARE (blue). Note that low- and high-SNR data were determined from unpaired time series. Data represent the average value of at least 3 measurements and shaded areas the respective standard deviation.*

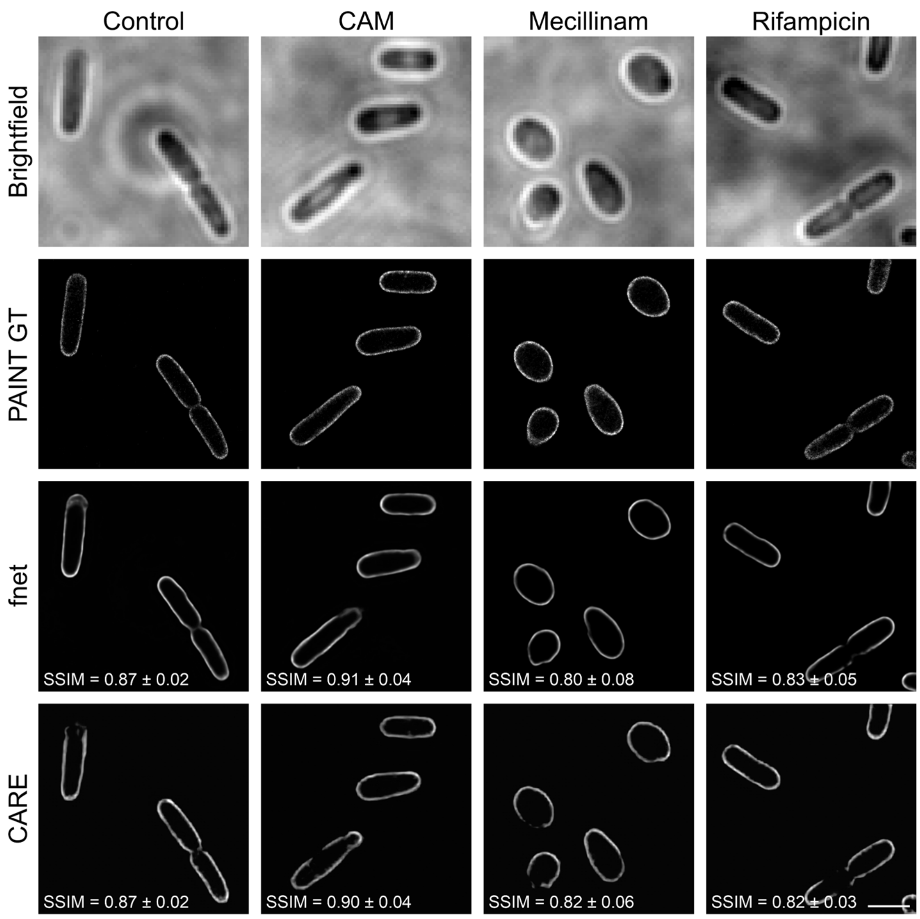


***Supplementary Figure 8:*** *Artificial labelling of E. coli membranes under different treatments. PAINT super-resolution images were used for model training. fnet and CARE models can also be used to predict membranes from cells that were treated with different antibiotics. SSIM = structural similarity. Scale bar is 2 µm.*

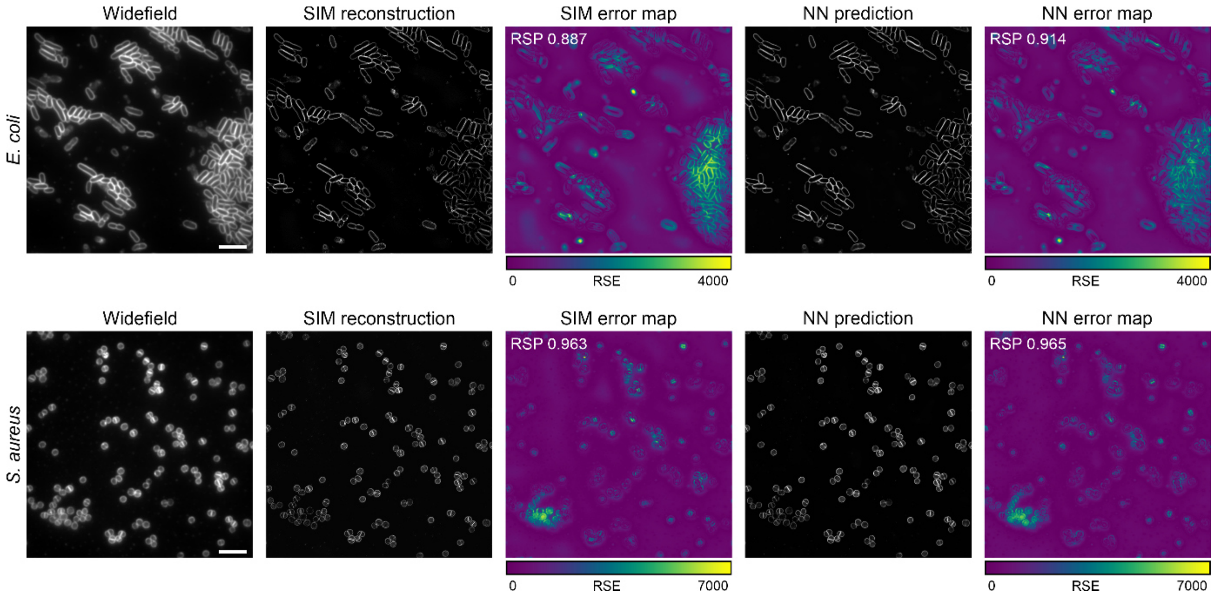


***Supplementary Figure 9****: SQUIRREL analysis* (Culley et al., 2018) *of SIM reconstructions and SIM images predicted by the NN (CARE). Error maps show the resolution scaled error (RSE) calculated between the blurred super-resolution image and the widefield image. The resolution-scaled Pearson coefficient (RSP) represents a normalised quality metric, providing high values and thus high-quality reconstructions for both SIM images and NN predictions. Scale bars are 5 µm.*

### Supplementary Tables

*Supplementary Table 1: Dataset sizes and imaging modalities of the dataset provided in this study. Different test datasets were provided for 3B (stitched/individual images). BF: Bright field; WF: Widefield; Fl.: Fluorescence; SR: Super-resolution; PAINT: Point accumulation for imaging in nanoscale topography; SIM: structured illumination microscopy; FoV: Field of view.*

| Figure | Task | Species | Training images | Test images | Image size (px) | Image size (µm) | ∅ objects/image | Imaging modality |
|---|---|---|---|---|---|---|---|---|
| 2B | Segmentation | *S. aureus* | 28 | 5 | 256 x 256 | 20.45 x 20.45 | 45 ± 34 | BF / Fl. (WF) |
| 2C | Segmentation | *E. coli* | 19 | 15 | 1024 x 1024 | 81.05 x 81.05 | 81 ± 24 | BF (WF) |
| 2D | Segmentation | *B. subtilis* | 80 | 10 | 1024 x 1024 | 66.56 x 66.56 | 53 ± 31 | Fl. (WF) |
| 3A | Object detection | *E. coli* | 25 | 15 | 512 x 512 | 81.05 x 81.05 | 52 ± 10 | BF (WF) (large FoV) |
| 3B | Object detection | *E. coli* | 153 | 32/50 | 400 x 400 | 33.6 x 33.6 | 15 ± 8 | Fl. (confocal) |
| 4A | Denoising | *E. coli* | 28 | 2 | 512 x 512 | 23.11 x 23.11 | 26 ± 19 | Fl. (confocal) |
| 4E | Denoising | *B. subtilis* | 3 | - | 1024 x 1024 | 66.56 x 66.56 | 110 ± 13 | Fl. (WF) |
| 5A | Artificial labelling | *E. coli* | 56 | 8 | 512 x 512 | 27.14 x 27.14 | 15 ± 7 | BF / Fl (WF) |
| 5A | Artificial labelling | *E. coli* | 33 | 20 | 512 x 512 | 10.01 x 10.01 | 1 - 4 | BF / SR (PAINT) |
| 6A | SR prediction | *E. coli* | 55 | 5 | 512 x 512 | 40.96 x 40.96 | ~ 50 - 300 | SR (SIM) |
| 6B | SR prediction | *S. aureus* | 94 | 5 | 512 x 512 | 40.96 x 40.96 | 96 ± 52 | SR (SIM) |

*Supplementary Table 2: List of training/test data and models shared via Zenodo*

| Figure | Task | Species | DOI | Type |
|---|---|---|---|---|
| 2B | Segmentation | *S. aureus* | 10.5281/zenodo.5550933 | Training/test data |
| 2C | Segmentation | *E. coli* | 10.5281/zenodo.5550935 | Training/test data |
| 2D | Segmentation | *B. subtilis* | 10.5281/zenodo.5550968 | Training/test data |
| 2D | Segmentation | *B. subtilis* | 10.5281/zenodo.5639253 | Training/test data + Multilabel-U-Net model |
| S2 | Segmentation | Mixed | 10.5281/zenodo.5551009 | Training/test data + StarDist 2D model |
| 3A | Object detection | *E. coli* | 10.5281/zenodo.5551016 | Training/test data + YOLOv2 model |
| 3B | Object detection | *E. coli* | 10.5281/zenodo.5551057 | Training/test data + YOLOv2 model |
| 4A | Denoising | *E. coli* | 10.5281/zenodo.5551112 | Training/test data + CARE 2D model |
| 4E | Denoising | *B. subtilis* | 10.5281/zenodo.5551135 | Training/test data |
| 5A | Artificial labelling | *E. coli* | 10.5281/zenodo.5551123 | Training/test data + fnet/CARE models |
| 6A | SR prediction | *E. coli* | 10.5281/zenodo.5551153 | Training/test data + CARE model |
| 6B | SR prediction | *S. aureus* | 10.5281/zenodo.5551141 | Training/test data + CARE model |

*Supplementary Table 3: Training parameters for segmentation networks*

| Organism | Mode | Network | Images (train/test) | Epochs | Steps | Image size | Patch size | Batch size | LR | % Valid. | Augmentation | Misc. | Train time |
|---|---|---|---|---|---|---|---|---|---|---|---|---|---|
| S. aureus | Brightfield | StarDist | 28/5 | 400 | 22 | 256 x 256 $px^2$ | 256 x 256 $px^2$ | 4 | 0.0003 | 20 | 4x (flip/rot.) | grid 1, 32 rays | 24 min 53 sec |
| | Fluorescence | StarDist | 28/5 | 400 | 24 | 256 x 256 $px^2$ | 256 x 256 $px^2$ | 4 | 0.0003 | 20 | 4x (flip/rot.) | grid 1, 32 rays | 53 min 27 sec |
| E. coli | Brightfield | U-Net | 19/15 | 100 | 30 | 1024 x 1024 $px^2$ | 512 x512 $px^2$ | 2 | 0.0003 | 10 | Flip/Shear | | 23 min |
| | | ML-U-Net | 19/15 | 500 | 17 | 1024 x 1024 $px^2$ | 512 x512 $px^2$ | 4 | 0.0003 | 10 | 4x (all options) | 2 pooling steps | 1 h 8 min |
| | | CARE | 19/15 | 100 | 300 | 1024 x 1024 $px^2$ | 128 x 128 $px^2$ | 8 | 0.0004 | 20 | 4x (flip/rot.) | | 42 min 16 sec |
| | | StarDist | 19/15 | 200 | 30 | 1024 x 1024 $px^2$ | 512 x512 $px^2$ | 2 | 0.0003 | 20 | 4x (flip/rot.) | grid 1, 80 rays | 45 min 23 sec |
| | | pix2pix | 19/15 | 500 | n.a. | 1024 x 1024 $px^2$ | 256 x 256 $px^2$ | 1 | 0.0002 | n.a. | 4x (flip/rot.) | | 47 min 57 sec |
| B. subtilis | Fluorescence | U-Net | 80/10 | 100 | 29 | 1024 x 1024 $px^2$ | 512 x 512 $px^2$ | 10 | 0.001 | 10 | 4x (flip/rot.) | | 47 min |
| | | ML-U-Net | 80/10 | 200 | 83 | 1024 x 1024 $px^2$ | 256 x 256 $px^2$ | 8 | 0.0005 | 10 | 4x (all options) | 3 pooling steps | 1 h 10 min |
| | | CARE | 80/10 | 50 | 2000 | 1024 x 1024 $px^2$ | 64 x 64 $px^2$ | 16 | 0.001 | 20 | 4x (flip/rot.) | | 1h 7 min |
| | | StarDist | 80/10 | 400 | 80 | 1024 x 1024 $px^2$ | 512 x 512 $px^2$ | 4 | 0.0003 | 10 | 4x (flip/rot.) | grid 2, 64 rays | 4 h 20 min |
| | | SplineDist | 32/10 | 800 | 25 | 512 x 512 $px^2$ | 256 x 256 $px^2$ | 4 | 0.0003 | 20 | 4x (flip/rot.) | grid 2 | 5 h 11 min |
| | | pix2pix | 80/10 | 1000 | n.a. | 1024 x 1024 $px^2$ | 256 x 256 $px^2$ | 1 | 0.0002 | n.a. | 4x (flip/rot.) | | 5h 50 min |
| All above | mixed model | StarDist | 155/35 | 200 | 120 | all above | 256 x 256 $px^2$ | 4 | 0.0003 | 10 | 3x (flip/rot.) | grid 2, 64 rays | 44 min 34 sec |

*Supplementary Table 4: Segmentation performance of StarDist generalist model (G) and specialist models (S)*

| Dataset | IoU (G) | IoU (S) | Precision (G) | Precision (S) | Recall (G) | Recall (S) |
|---|---|---|---|---|---|---|
| *E. coli* bright field | 0.74 ± 0.03 | 0.78 ± 0.03 | 0.88 ± 0.08 | 0.87 ± 0.07 | 0.82 ± 0.12 | 0.83 ± 0.12 |
| *S. aureus* bright field | 0.71 ± 0.02 | 0.64 ± 0.01 | 0.89 ± 0.04 | 0.90 ± 0.02 | 0.99 ± 0.02 | 0.87 ± 0.03 |
| *S. aureus* fluorescence | 0.86 ± 0.01 | 0.91 ± 0.03 | 0.95 ± 0.03 | 0.98 ± 0.02 | 0.99 ± 0.02 | 1.00 ± 0.01 |
| *B. subtilis* fluorescence | 0.69 ± 0.04 | 0.76 ± 0.03 | 0.82 ± 0.09 | 0.92 ± 0.04 | 0.73 ± 0.09 | 0.88 ± 0.55 |

*Supplementary Table 5: YOLOv2 model training parameters*

| Dataset | Images (train/test) | Epochs | Image size | Train cycles | Batch size | LR | % Valid. | Augmentation | Penalties (FNP, FPP, PSP, FCP) | Training time |
|---|---|---|---|---|---|---|---|---|---|---|
| Growth stage (large FoV) | 25/15 | 100 | 512 x 512 px² | 4 | 4 | 0.0003 | 20 | 4x (flip/rot.) | 5, 1, 3, 3 | 1 h 31 min |
| Growth stage (small FoV) | 100/60 | 100 | 256 x 256 px² | 4 | 8 | 0.0003 | 20 | 4x (flip/rot.) | 5, 1, 3, 3 | 34 min |
| Antibiotic profiling | 153/32 (50) | 100 | 400 x 400 px² | 4 | 16 | 0.001 | 20 | 8x (flip/rot.) | 4, 2, 1, 2 | 2 h 33 min |

*Supplementary Table 6: Performance of YOLOv2 models for growth stage prediction, trained on differently sized field of views (FOV). mean average precision (mAP) values were calculated from the individual classes' average precision (AP) values.*

| FOV | Class | Recall | Precision | AP | mAP |
|---|---|---|---|---|---|
| Large (512 x 512 px²) | Rod | 0.442 | 0.621 | 0.321 | **0.386** |
| | Dividing | 0.414 | 0.760 | 0.358 | |
| | Microcolony | 0.567 | 0.567 | 0.479 | |
| Small (256 x 256 px²) | Rod | 0.746 | 0.712 | 0.643 | **0.667** |
| | Dividing | 0.657 | 0.722 | 0.589 | |
| | Microcolony | 0.804 | 0.755 | 0.771 | |

*Supplementary Table 7: Supplier information and experimental conditions for the antibiotics used in this study*

| Antibiotic | Supplier | Cat. # | Working concentration | Exposure time |
|---|---|---|---|---|
| MP265 | Alfa Aesar | B25362 | 25 µM | 60 min |
| Mecillinam | Sigma Aldrich | 33447 | 2 µg/ml | 60 min |
| Rifampicin | Sigma Aldrich | R3501 | 100 µg/ml | 60 min |
| Chloramphenicol | Sigma Aldrich | C0378 | 50 µg/ml | 60 min |
| Nalidixate | Alfa Aesar | J63550 | 50 µg/ml | 30 min |

*Supplementary Table 8: Performance of the YOLOv2 antibiotic profiling model on different types of test datasets.*

| Test data | Class | # of objects in training dataset | # of objects in test dataset | Recall | Precision | AP score | mAP |
|---|---|---|---|---|---|---|---|
| stitched images | Rifampicin | 430 | 102 | 0.920 | 0.928 | 0.879 | **0.662** |
| | Oblique | 535 | 93 | 0.871 | 0.750 | 0.769 | |
| | Nalidixate | 87 | 58 | 0.707 | 0.854 | 0.641 | |
| | Mecillinam | 327 | 91 | 0.703 | 0.744 | 0.605 | |
| | CAM | 264 | 45 | 0.978 | 0.620 | 0.754 | |
| | Vesicle | 207 | 67 | 0.284 | 0.594 | 0.207 | |
| | Control | 181 | 66 | 0.939 | 0.912 | 0.914 | |
| | MP265 | 265 | 78 | 0.654 | 0.671 | 0.526 | |
| individual treatment | Rifampicin | 430 | 55 | 0.891 | 0.907 | 0.860 | **0.690** |
| | Oblique | 535 | 161 | 0.745 | 0.876 | 0.713 | |
| | Nalidixate | 87 | 64 | 0.672 | 0.796 | 0.602 | |
| | Mecillinam | 327 | 66 | 0.758 | 0.769 | 0.694 | |
| | CAM | 264 | 65 | 0.938 | 0.635 | 0.759 | |
| | Vesicle | 207 | 119 | 0.353 | 0.600 | 0.257 | |
| | Control | 181 | 61 | 0.984 | 0.822 | 0.944 | |
| | MP265 | 265 | 76 | 0.789 | 0.800 | 0.692 | |

*Supplementary Table 9: Model training parameters for image denoising*

| Organism | Network | Images (train/test) | Epochs | Steps | Image size | Patch size (#/img) | Batch size | LR | % Valid. | Augmentation | Train time |
|---|---|---|---|---|---|---|---|---|---|---|---|
| E. coli | CARE | 28/2 | 100 | 600 | 512 x 512 $px^2$ | 64 x 64 $px^2$ (50) | 8 | 0.0004 | 10 | 4x (flip/rot.) | 30 min 34 s |
| | N2V | 28/2 | 200 | 101 | 512 x 512 $px^2$ | 64 x 64 $px^2$ (50) | 128 | 0.0004 | 10 | 4x (flip/rot.) | 41 min |
| B. subtilis | N2V | 3/n.a. | 200 | 44 | 1024 x 1024 $px^2$ | 64 x 64 $px^2$ (500) | 128 | 0.0004 | 10 | 4x (flip/rot.) | 18 min |

*Supplementary Table 10: Model training parameters for artificial labelling*

| Imaging Mode | Scaling | Network | Images (train/test) | Epochs | Steps | Image size | Patch size (#/img) | Batch size | LR | % Valid. | Augmentation | Train time |
|---|---|---|---|---|---|---|---|---|---|---|---|---|
| Widefield | none | fnet | 56/8 | n.a. | 200000 | 256 x 256 $px^2$ | 128 x 128 $px^2$ (1) | 2 | 0.001 | 10 | 4x (flip/rot.) | 1 h 43 min |
| (108 + 158 nm/px) | | CARE | 56/8 | 300 | 200 | 256 x 256 $px^2$ | 64 x 64 $px^2$ (4) | 4 | 0.0004 | 10 | 4x (flip/rot.) | 57 min |
| PAINT | 8x | fnet | 33/20 | n.a. | 200000 | 512 x 512 $px^2$ | 128 x 128 $px^2$ (1) | 4 | 0.0004 | 10 | 4x (flip/rot.) | 2 h 27 min |
| (158 nm/px) | | CARE | 33/20 | 300 | 100 | 512 x 512 $px^2$ | 256 x 256 $px^2$ (1) | 4 | 0.0004 | 10 | 4x (flip/rot.) | 1 h 33 min |

*Supplementary Table 11: Model training parameters for super-resolution prediction*

| Organism | Scaling | Network | Images (train/test) | Epochs | Steps | Image size | Patch size (#/img) | Batch size | LR | % Valid. | Augmentation | Train time |
|---|---|---|---|---|---|---|---|---|---|---|---|---|
| E. coli | 2x | CARE | 55/5 (link) | 300 | 2500 | 1024 x 1024 $px^2$ | 80 x 80 $px^2$ (100) | 8 | 0.0004 | 10 | 4x (flip/rot.) | 6 h 6 min |
| S. aureus | 2x | CARE | 94/5 (link) | 300 | 3000 | 1024 x 1024 $px^2$ | 80 x 80 $px^2$ (100) | 8 | 0.0004 | 10 | 4x (flip/rot.) | 7 h 23 min |

#### Supplementary Videos:

*Supplementary Video 1: Application of a StarDist model trained using the ZerCostDL4Mic platform in the StarDist Fiji plugin.*

*Supplementary Video 2: StarDist segmentation of S. aureus time-lapse data at high cell density. Yellow lines outline segmented cells.*

*Supplementary Video 3: Segmentation and tracking of E. coli time-lapse data using a multilabel-U-Net and TrackMate. Boundaries are coloured according to the lineages.*

*Supplementary Video 4: YOLOv2 object detection of different growth stages in E. coli time-lapse data. The model was trained to detect non-dividing cells (blue bounding boxes), dividing cells (green bounding boxes) or microcolonies (red bounding boxes) using the entire field of view (80x80 µm²).*

*Supplementary Video 5: YOLOv2 object detection of different growth stages in E. coli time-lapse data. The model was trained to detect non-dividing cells (blue bounding boxes), dividing cells (green bounding boxes) or microcolonies (4+ cells in close contact, red bounding boxes) using a ROI of 40x40 µm².*

*Supplementary Video 6: Denoising of E. coli nucleoid dynamics. Confocal time-lapse videos of E. coli cells expressing H-NS-mScarlet protein fusion from the native locus were denoised using PureDenoise, Noise2Void or CARE.*

*Supplementary Video 7: Self-supervised denoising of FtsZ-GFP dynamics in vertically aligned B. subtilis cells using Noise2Void. The false-coloured panel is shown for visualisation. Scale bar is 1 µm.*

*Supplementary Video 8: Artificial labelling of E. coli bright field time series using fnet. The fnet 2D model was trained using paired bright field and super-resolution PAINT images.*

*Supplementary Video 9: Prediction of super-resolution SIM images from widefield fluorescence images. Live S. aureus cells were stained with Nile Red. The right panel shows the overlay between SIM reconstructions (green) and the CARE 2D prediction (magenta).*

#### *References*


Chamier, L. von, Laine, R.F., Jukkala, J., Spahn, C., Krentzel, D., Nehme, E., *et al.* (2021) Democratising deep learning for microscopy with ZeroCostDL4Mic. *Nat Commun* **12**: 2276.

Culley, S., Albrecht, D., Jacobs, C., Pereira, P.M., Leterrier, C., Mercer, J., and Henriques, R. (2018) Quantitative mapping and minimization of super-resolution optical imaging artifacts. *Nat Methods* **15**: 263–266.

Kass, M., Witkin, A., and Terzopoulos, D. (1988) Snakes: Active contour models. *International Journal of Computer Vision* **1**: 321–331 http://dx.doi.org/10.1007/bf00133570.

Paintdakhi, A., Parry, B., Campos, M., Irnov, I., Elf, J., Surovtsev, I., and Jacobs-Wagner, C. (2016) Oufti: an integrated software package for high-accuracy, high-throughput quantitative microscopy analysis. *Mol Microbiol* **99**: 767–777.

Panigrahi, S., Murat, D., Le Gall, A., Martineau, E., Goldlust, K., Fiche, J.-B., *et al.* (2021) Misic, a general deep learning-based method for the high-throughput cell segmentation of complex bacterial communities. https://elifesciences.org/articles/65151. Accessed October 5, 2021.

Stylianidou, S., Brennan, C., Nissen, S.B., Kuwada, N.J., and Wiggins, P.A. (2016) SuperSegger: robust image segmentation, analysis and lineage tracking of bacterial cells. *Mol Microbiol* **102**: 690–700.

Van Valen, D.A., Kudo, T., Lane, K.M., Macklin, D.N., Quach, N.T., DeFelice, M.M., *et al.* (2016) Deep Learning Automates the Quantitative Analysis of Individual Cells in Live-Cell Imaging Experiments. *PLoS Comput Biol* **12**: e1005177.

Wang, Z., Bovik, A.C., Sheikh, H.R., and Simoncelli, E.P. (2004) Image quality assessment: from error visibility to structural similarity. *IEEE Trans Image Process* **13**: 600–612.
